# A Systematic Assessment of Robustness in CNS Safety Pharmacology

**DOI:** 10.1101/2024.03.21.586096

**Authors:** Maria Reiber, Helen Stirling, Tim P. Ahuis, Washington Arias, Katharina Aulehner, Ute Dreßler, Martien J.H. Kas, Johanna Kela, Kimberly Kerker, Tarja Kuosmanen, Helga Lorenz, Alexander T. Pennington, Eva-Lotta von Rüden, Heike Schauerte, Isabel Seiffert, Steven R. Talbot, Christina Torturo, Sami Virtanen, Ann-Marie Waldron, Sylvie Ramboz, Heidrun Potschka

## Abstract

Irwin tests are key preclinical study elements for characterizing drug-induced neurological side effects. This multicenter study aimed to assess the robustness of Irwin tests across multinational sites during three stages of protocol harmonization. The projects were part of the EQIPD framework (<u>E</u>nhanced <u>Q</u>uality in <u>P</u>reclinical <u>D</u>ata, https://quality-preclinical-data.eu/), aiming to increase success rates in transition from preclinical testing to clinical application. Female and male NMRI mice were assigned to one of three groups (vehicle, 0.1 mg/kg MK-801, 0.3 mg/kg MK-801). Irwin scores were assessed at baseline and multiple times following injection of MK-801, a non-competitive NMDA antagonist, using local protocols (stage 1), a shared protocol with harmonized environmental design (stage 2), and fully harmonized Irwin scoring protocols (stage 3). The analysis based on the four functional domains (motor, autonomic, sedation, and excitation) revealed substantial data variability in stages 1 and 2. Although there was still marked overall heterogeneity between sites in stage 3 after complete harmonization of the Irwin scoring scheme, heterogeneity was only moderate within functional domains. When comparing treatment groups vs. vehicle, we found large effect sizes in the motor domain and subtle to moderate effects in the excitation-related and autonomic domain. The pronounced interlaboratory variability in Irwin datasets for the CNS-active compound MK-801 needs to be carefully considered by companies and experimenters when making decisions during drug development. While environmental and general study design had a minor impact, the study suggests that harmonization of parameters and their scoring can limit variability and increase robustness.

## Introduction

Limited replicability of results is discussed as one of the major obstacles in preclinical animal-based research and its translation to the clinic [e.g., (1-5)]. For decades, it was assumed that harmonization and standardization of experimental procedures would significantly contribute to the successful translation of preclinical testing to clinical application. However, doubts have arisen about the infallibility of standardization as a means of addressing poor replicability in preclinical study designs (3, 6-8).

The multinational EQIPD consortium (<u>E</u>nhanced <u>Q</u>uality in <u>P</u>reclinical <u>D</u>ata, https://quality-preclinical-data.eu/) aimed to identify factors influencing the quality of data generated in preclinical research as a basis for recommendations enabling a smoother and successful transition from preclinical research to clinical application (9, 10). In a recent study, the consortium analyzed the impact of protocol harmonization on the replicability of data in the open field test, i.e., a behavioral paradigm with automated recording of the primary outcome parameters (11). The study demonstrated that harmonization of protocols can reduce between-site variability (11). Considering that the standardized application of scoring systems and their harmonization across sites can pose a particular challenge, we next addressed the question of whether a comparable impact can be observed for a frequently used paradigm, based on scoring of a variety of readout parameters by experimenters.

As introduced by Samuel Irwin (1968), the Irwin test is a comprehensive observational assessment, i.e., a systematic, quantitative procedure for measuring mice’s behavioral and physiological state and their response to drugs (12). Thereby, different parameters are evaluated based on scoring systems to assess and grade the effects of drug candidates on the central, peripheral, and autonomic nervous system. Since its conception, various modified Irwin test versions have been designed considering specific aims and needs in drug development [e.g., (13, 14)]. Moreover, cross-species test protocols for comparable functional observation batteries (FOBs) have been successfully validated for use in rats, dogs, non-human primates, and mini-pigs [(15-21)], expanding its scope of application. Irwin and other FOBs are primarily used for safety assessment in CNS drug development, providing information for decision-making, candidate selection, or the need for more in-depth analysis of selected potential adverse effects (22). According to the ICH S7A document ‘*Safety pharmacology studies for human pharmaceuticals – scientific guideline*’, these tests are considered key preclinical study elements that are prerequisites for proceeding to clinical trials (23). Irwin scoring and interpreting of animals’ behaviors go hand-in-hand with direct interaction between the examiner and the animal. Therefore, thorough education and adequate training of test examiners is crucial. A recent study has evaluated the impact of partial standardization focussed on a standard dose, administration route, and time points on Irwin/FOB data obtained by testing two drugs in male Wistar rats based on laboratory-specific scoring protocols (24). The authors reported that their partial standardization did not result in a relevant reduction in quantitative variability.

In this study, we used a comparable test scenario as a starting point with the same dose, route of administration, and timeline. The impact of further stepwise harmonization of scoring protocols across sites was then assessed. The main aim was to compare harmonized protocols with original heterogeneous versions across sites to determine whether the harmonization procedure affects variability, replicability, robustness, and/or external validity of results. Therefore, we analyzed the influence of protocols in three experimental stages based on the application of 1. local protocols without harmonization, 2. a shared protocol with harmonized experimental and environmental factors, and 3. a shared protocol with harmonized experimental (including identical scoring schemes) and environmental factors.

To allow comparability with two other subprojects of the EQIPD consortium that focused on an open field paradigm (11) and pharmaco-EEG, the study was performed in mice, representing the second most common species for Irwin/FOB testing (16, 24). MK-801 was chosen as the compound, considering its broad range of effects on various parameters from different functional domains assessed in the Irwin test.

## Results

The basis for evaluating Irwin data in a multicenter approach was comparing laboratory scores. Because the Irwin score consists of many parameters, we subclassified parameters. We designed main observational groups (functional domains) that may reflect the multidimensional aspect of the Irwin/FOB test. With this approach, we aimed to avoid bias that may arise from the selection and analysis of individual parameters. Therefore, outcome measures were split into the following four functional domains. First, the motor functions-related domain (‘motor domain’), including coordination, gait, and muscle tone; second, the autonomic functions-related domain (‘autonomic domain’); third, the excitation-related domain; fourth, the sedation-related domain. Individual parameters that could not be assigned to these functional domains were classified as ‘other measures’, comprising Straub tail, toe pinch/ tail pinch reflex, and writhing. ‘Other measures’ readouts are presented in **Appendix 1**.

The individual scoring parameters assigned to the four domains differ between the sites in stages 1 and 2 because the scoring system was not harmonized before stage 3. An overview of the site-specific subsummation of parameters is provided in **Appendix 2**. If necessary, variables were recoded so that an increase and decrease contributed comparably to the sum score. Adherence to this procedure was required for the following two variables: fearfulness and touch response. Moreover, the following variables were split into two domains depending on whether a score was increased or decreased: alertness and irritability, i.e., increase contributed to the excitation-related domain, whereas decrease contributed to the sedation-related domain. Because the directionality of the startle response was not provided from enough sites, this variable could not be included in the analyses.

### Stages 1 and 2

In stage 1, each of the 5 sites followed its in-house SOP for Irwin testing except for the basic specifications described above (e.g., experimental cohorts, strain, and compound). An overview of the results, illustrated per site and domain as change-from-baseline scores, is presented in **Figure 1**. Stage 1 results of the parameters subsumed under ‘other measures’ are presented in **S1, S2, S3, S4, and S5 Supplementary Figures**.

**Figure 1.**
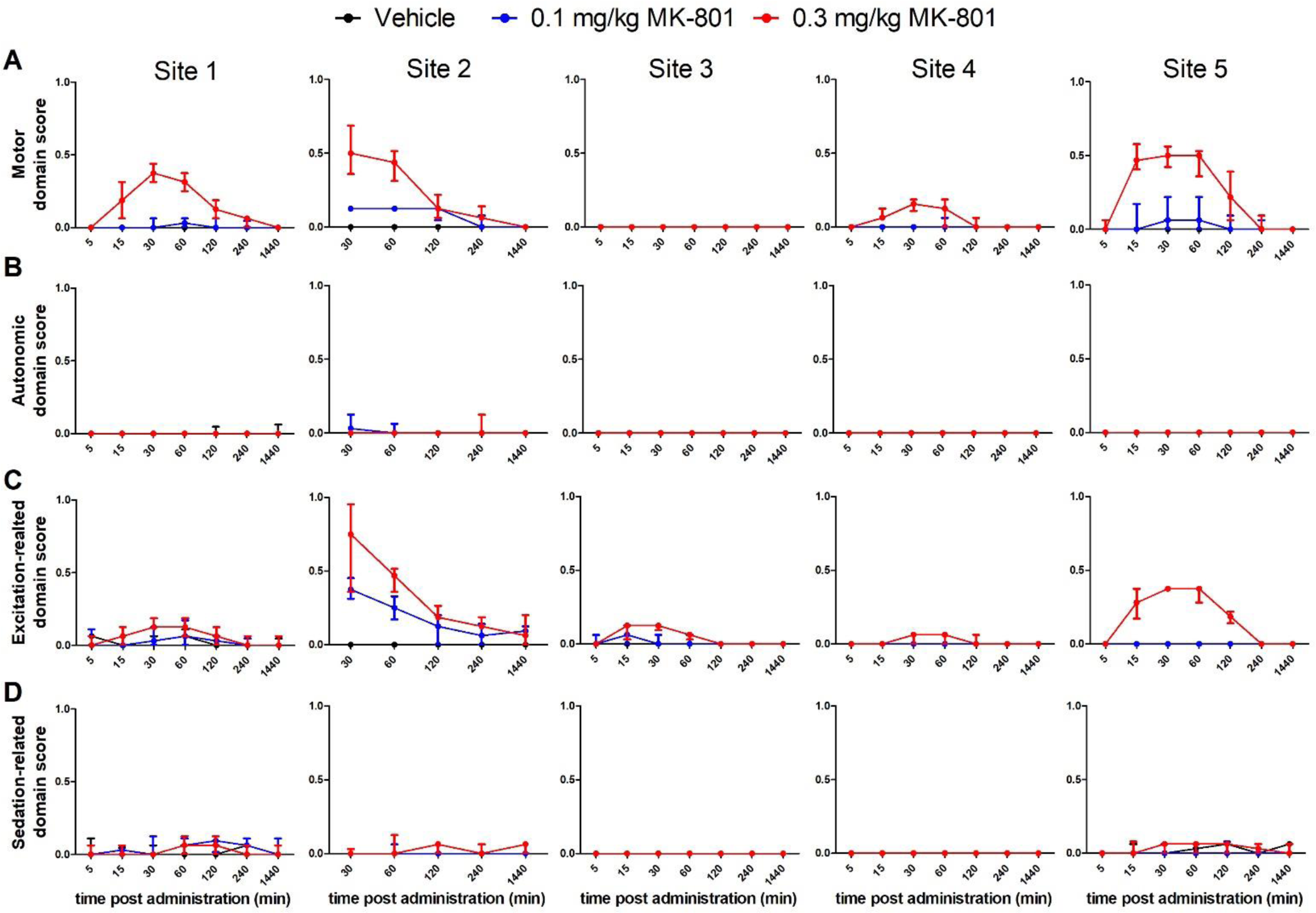
Visualization of stage 1 data across laboratory sites. Delta sum scores (positive change-from-baseline scores) for each functional domain are presented as median with interquartile range (IQR). Site-specific Irwin outcome measures were assigned to the following functional domains for centralized analysis: **(A)** motor, **(B)** autonomic, **(C)** excitation-related, **(D)** sedation-related domain. Details about the site-specific scoring schemes are provided in **S1 and S2** (Site 1)**, S3 and S4** (Site 2), **S5 and S6** (Site 3), **S7 and S8** (Site 4), **S9 and S10** (Site 5) **Supplementary Tables**. *n* = 11-12 per group (site 1); *n* = 10 per group (site 2, site 4, and site 5); *n* = 9-11 per group (site 3). The datasets underlying this figure are available in the Figshare Repository [DOI will be provided in the accepted article].

In stage 2, critical variables concerning the environmental and experimental setting were harmonized across the 5 laboratory sites. **Figure 2** illustrates change-from-baseline scores per domain for stage 2. Stage 2 results of the parameters subsumed under ‘other measures’ are presented in **S6, S7, S8, S9, and S10 Supplementary Figures**.

**Figure 2.**
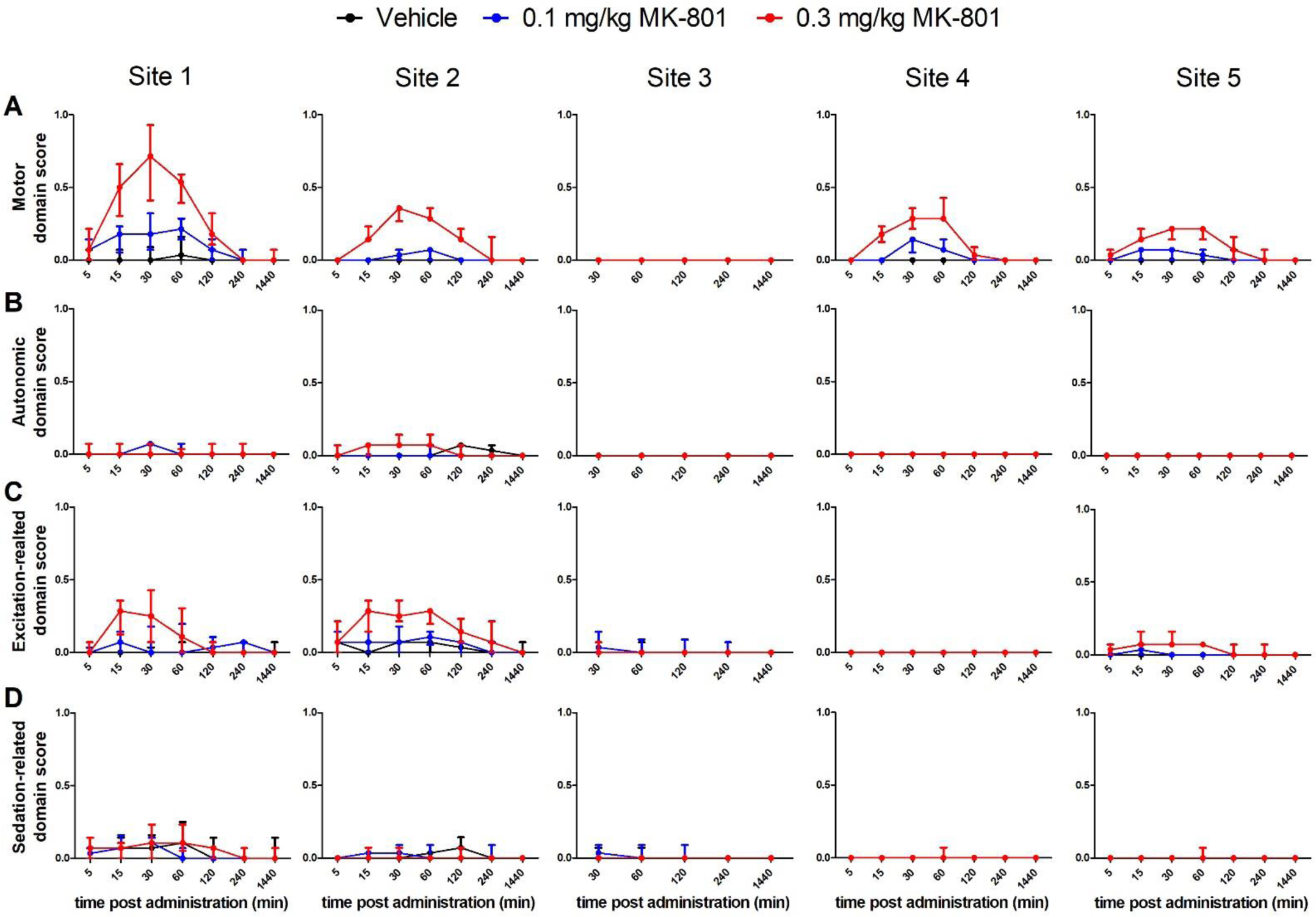
Visualization of stage 2 data across laboratory sites. Delta sum scores (positive change-from-baseline scores) for each functional domain are presented as median with interquartile range (IQR). Site-specific Irwin outcome measures were assigned to the following functional domains for centralized analysis: **(A)** motor, **(B)** autonomic, **(C)** excitation-related, **(D)** sedation-related domain. Details about the site-specific scoring schemes are provided in **S19 Supplementary Table**. *n* = 10 per group per site. The datasets underlying this figure are available in the Figshare Repository [DOI will be provided in the accepted article].

Overall, delta score illustrations suggested that MK-801-dependent effects were most pronounced across sites in the motor and excitation-related domains. Moreover, stage 2 plots indicated that motor-domain delta scores reflected dose-dependent effects in four out of five labs: the scoring parameters allocated to the motor domain also displayed the effects of the lower dosage. Interestingly, one lab did not measure relevant effects of MK-801 in any of the four functional domains using its in-house SOP in stages 1 and 2.

Due to the heterogeneity of the outcome measures in stages 1 and 2 based on site-specific scoring schemes, we have not conducted analyses to compare effect sizes across sites.

### Stage 3

For stage 3, a fully harmonized scoring system was applied across the three participating sites. A domain-specific overview of the change-from-baseline scores of the data from the three participating sites is provided in **Figure 3**. The results of the parameters subsumed under ‘other measures’ are presented in **S11 Supplementary Figure**.

**Figure 3.**
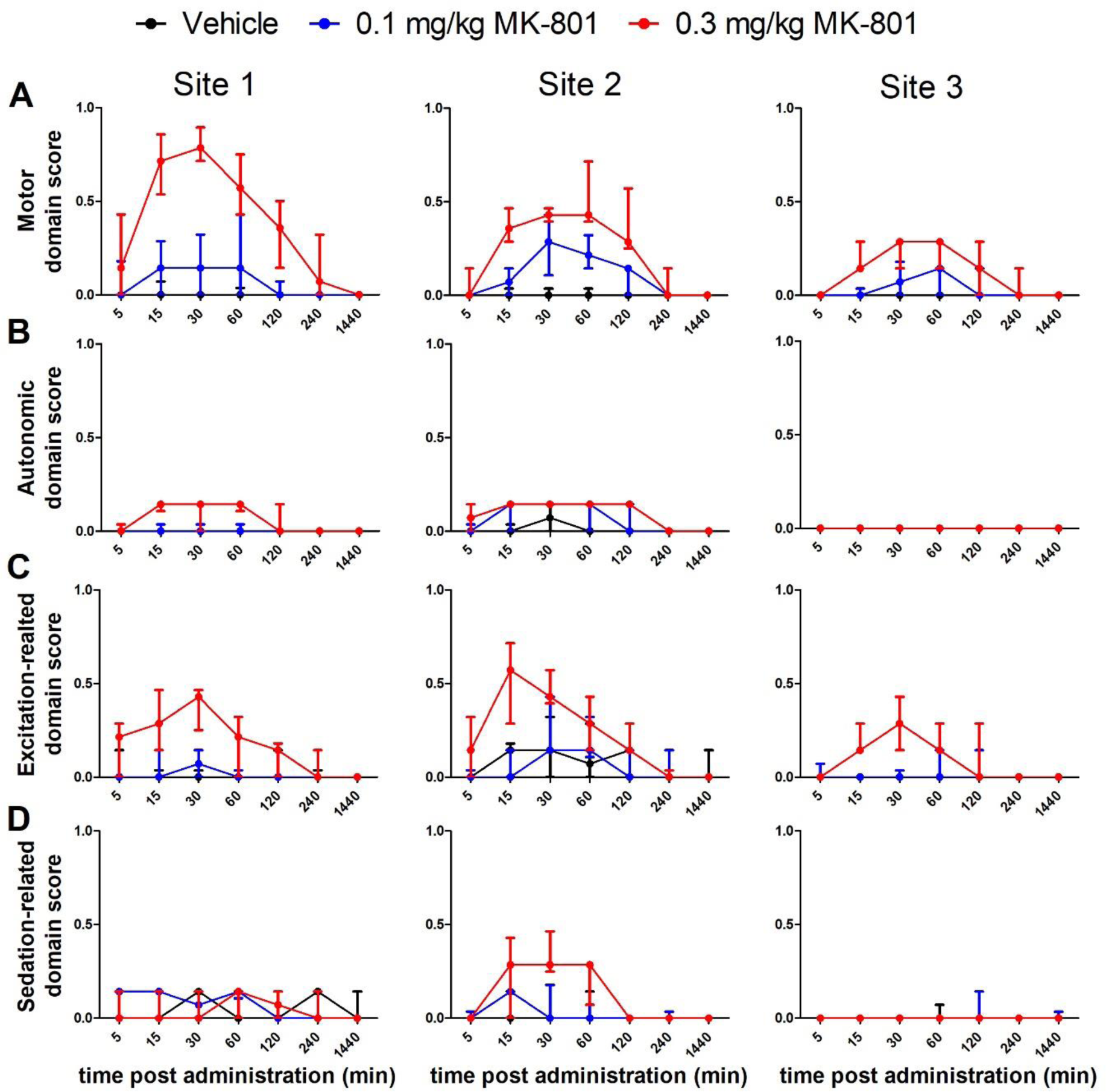
Visualization of stage 3 data across laboratory sites. Delta sum scores (positive change-from-baseline scores) for each functional domain are presented as median with interquartile range (IQR). The ring-test protocol used in stage 3 provided a fully harmonized Irwin scoring scheme. Irwin outcome measures were assigned to the following functional domains for centralized analysis: **(A)** motor, **(B)** autonomic, **(C)** excitation-related, **(D)** sedation-related domain. Details about the scoring scheme provided by the shared protocol can be found in **S20 Supplementary Table**. *n* = 10 per group (site 1 and site 2); *n* = 9-11 per group (site 3). The datasets underlying this figure are available in the Figshare Repository [DOI will be provided in the accepted article].

As laboratory 4 participated in stage 3, but did not follow the shared protocol for a harmonized scoring scheme and did not collect baseline data, records from this site were excluded from the centralized analysis. These data are provided as sum scores in **S12 and S13 Supplementary Figures**.

With respect to the centralized analysis, the illustration of the change-from-baseline scores revealed that the strongest MK-801-associated effects were found 30 minutes post-injection. Moreover, change-from-baseline scores suggested that dose-dependent effects displayed in the motor domain were more pronounced in stage 3 than in stages 1 and 2 before scoring-scheme harmonization. In the motor domain, no significant effect was found in the vehicle group across the three stages, which may indicate that findings for the motor domain were relatively robust.

A meta-analysis assessed the effect sizes of the two treatment groups (0.1 mg/kg MK-801 vs. 0.3 mg/kg MK-801) 30 minutes post-injection across labs. The results are presented per domain in a forest plot (**Figure 4**).

**Figure 4.**
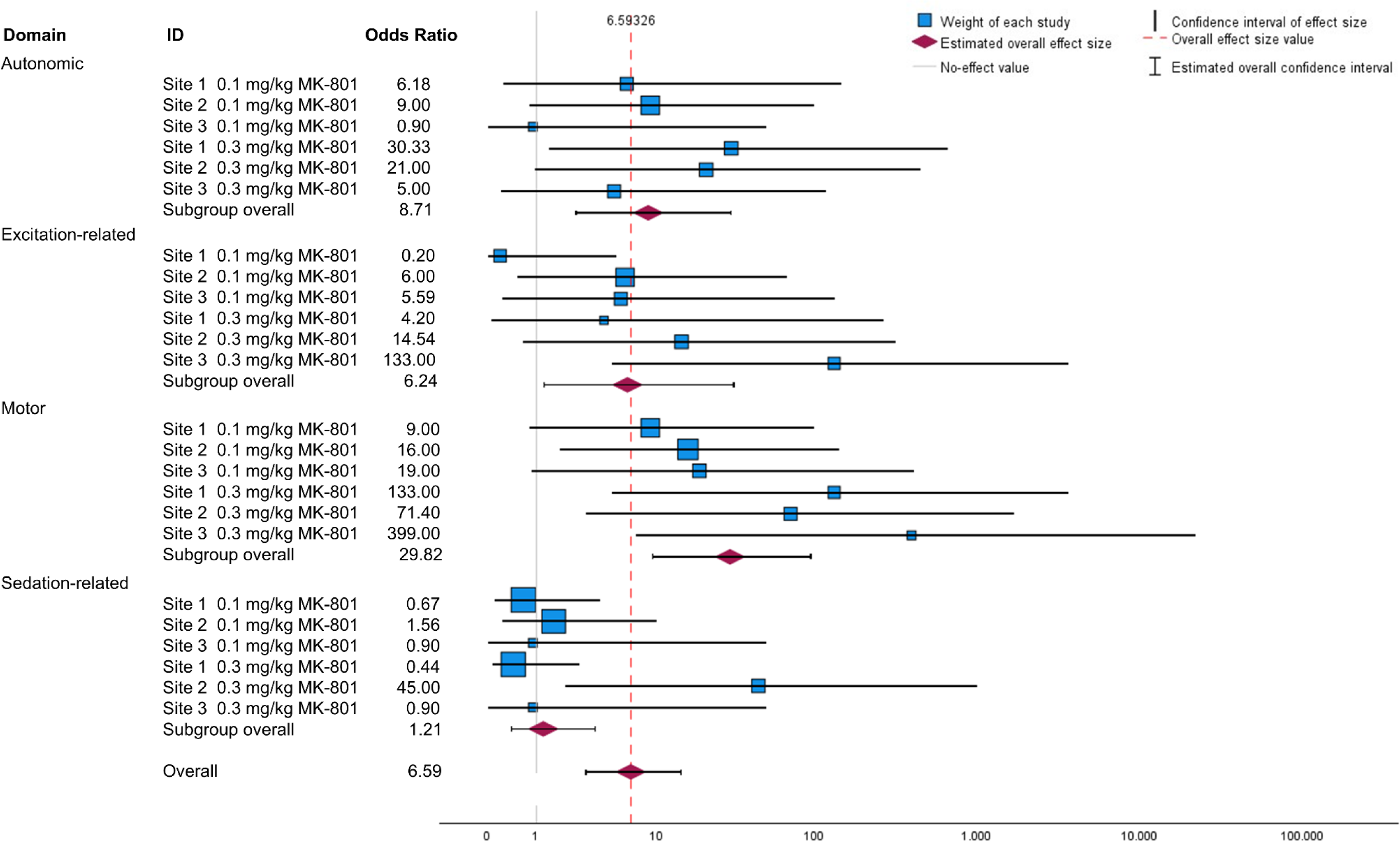
Forest plot of the meta-analysis conducted for stage 3 data. Domain-specific delta sum scores (positive change-from-baseline scores) 30 minutes post-injection were recoded on a binary scale to calculate effect sizes (Odds Ratios). The datasets underlying this figure are available in the Figshare Repository [DOI will be provided in the accepted article].

Overall, moderate heterogeneity was observed (I^2^ = 43,4%). Moderate heterogeneity within the functional domains was corroborated by Chi-Square tests not rejecting the H0 hypothesis of no relevant heterogeneity in the motor (*X*^2^ (1, *N* = 5) = 4.015, p = 0.55), autonomic (*X*^2^ (1, *N* = 5) = 2.330, p = 0.80), excitation-related (*X*^2^ (1, *N* = 5) = 7.877, p = 0.16), and sedation-related domain (*X*^2^ (1, *N* = 5) = 6.922, p = 0.23). However, the overlap of the included studies’ point estimates and their 95% CIs in the visual display may represent substantial heterogeneity among all included studies in stage 3. In addition, the Chi-Square test across functional domains pointed to relevant heterogeneity within all included studies in stage 3 (*X*^2^ (1, *N* = 23) = 40.980, p = 0.01).

Moreover, the Chi Square homogeneity test across the four functional domains was significant (*X*^2^ (1, *N* = 3) = 15.973, p = 0.001), suggesting the suitability of the four functional domains as moderator variables for subsequent subgroup analysis.

The overall pooled effects from the included studies within the autonomic and motor domain were significantly larger than the overall effect among all included studies (p < 0.01, respectively). The overall pooled effect from the included studies of the excitation-related domain was significantly smaller than the overall effect among all included studies (p = 0.03). The overall pooled effect from the included studies of the sedation-related domain indicated no significant domain-specific effect in comparison to the overall effect among all included studies (p = 0.73). Overall, when comparing treatment groups vs. vehicle, we found large effect sizes in the excitation-related, autonomic, and motor domain, with the latter showing by far the largest effect size. Notably, the effect size in the sedation-based domain was significantly lower than in the other domains.

## Discussion

In this multicenter study, we assessed the impact of different levels of protocol harmonization on the robustness of Irwin scoring. It has already been emphasized that variability observed in Irwin and FOB testing may question the overall validity of information obtained in preclinical rodent safety assessments (24-26). In particular, the FOB/Irwin test used, the applied scoring system, and the level of experience and/or training of investigators can profoundly affect the comparability of study results (24). Also, environmental and experimental factors have been considered key factors confounding the robustness of FOB/Irwin tests (24, 27, 28). Therefore, in this 3-stage multinational approach, we assessed the impact of two further harmonization steps, starting from stage 1, which was characterized by a level of harmonization comparable to the study by Himmel and colleagues (2019) (24).

In the first stage, the five participating sites used scoring schemes based on their in-house SOPs. This implies that the parameters themselves, the number of parameters, the scaling of the parameters, and the scoring method were not harmonized. Notably, the scoring schemes also provided a varying range with respect to outcome measures. In this context, it is emphasized that variants of Irwin scores and FOBs are often developed based on a rational approach. With development of specific scoring schemes, experimenters can better consider indications relevant to the tested compounds, and, more importantly, the specific needs or vulnerabilities of the respective patient populations (e.g., ataxia is regarded more relevant for drugs developed for a geriatric patient population).

The marked overall heterogeneity of stage 1 data can probably be partly attributed to the different scoring systems of the companies and institutes.

Regardless of indication area and patient population, the question whether a broad scoring range provides an informative added value arises. If a wide scoring range is used, it must be described in detail. Adequate and comprehensive training of the staff performing the tests must be ensured in order to achieve a sufficient test-retest and inter-rater reliability. It should be considered that it is challenging to implement a broad scale in a standardized manner even within one laboratory with multiple investigators. As already emphasized by Himmel and colleagues (2019), greater efforts are needed to standardize the often subjective variables and the often examiner-dependent scoring to reduce variability (24). Using Himmel and colleagues’ findings as a starting point in stage 1 (24), we applied two further steps of protocol harmonization. In stage 2, however, we did not detect any relevant reduction in variability due to the harmonization of environment and test conditions, and we did not find a relevant reduction in variability until the scoring system was harmonized in stage 3. These findings indicate that the factors harmonized for stage 2 testing, concerning housing conditions, group allocation procedure, personnel, and experimental set-up, had no relevant impact on the overall variability of the Irwin tests. In this context, it is emphasized that a high variability with site-specific differences was pronounced regarding the presence and magnitude of specific drug effects, the time course with time of peak effect and duration of effects, and the dose dependence of the effects. These findings further support the observations in earlier studies, which reported a considerable interlaboratory variability for Irwin/FOBs (24-26).

Stage 3 data suggest that harmonization of the actual scoring system can limit variability. However, in this context it needs to be considered that only three laboratories contributed to this experimental phase and that there is still evidence for relevant discrepancies and interlaboratory variability, which can affect decision-making during drug development. This variability is likely related to the subjective scoring which implies poor interrater reliability. Moser and colleagues (1997) have already assessed the impact of collective training (26). Despite the coordinated and harmonized training, the authors reported relevant dataset variability. Automated analysis would resolve the issue of examiner bias. However, as already discussed by Himmel and colleagues (2019), fully automated Irwin scoring is not feasible for the full range of Irwin parameters (e.g., reflex testing) (26). In stage 3, the scoring results were recoded to perform the binary meta-analysis of delta scores. Due to the binary scale, information about the extent of the effects was lost. Reducing the scale to a binary outcome entails reducing informative value. However, for some parameters it may be of particular relevance to which extent an effect exists (e.g., mild vs. pronounced ataxia).

Both historical and experimental data analysis by Himmel and colleagues (2019) pointed to differences in variability for different Irwin/FOB parameters (24). Whereas activity, posture, muscle tone, stereotypies, startle reflex, and body temperature proved more robust, piloerection, palpebral closure, and respiration were among the less robust parameters. These findings align with our datasets for the different functional domains. In stage 3, the findings for the motor domain parameters proved to be relatively robust including a consistent demonstration of dose-dependent effects. However, in this context, it is notable that not all laboratories recorded relevant motor effects during stage 1 and 2 testing. While findings for excitation-related parameters were rather consistent in stage 3 experiments, data for the autonomic domain and sedation-related domain were characterized by a relatively high variability. Moreover, it needs to be considered that only subtle changes became evident for these parameters even in laboratories reporting respective effects. Thus, it remains difficult to compare the robustness of the different functional domains. In this context, it would be of future interest to conduct a multicenter study with compounds that exert more pronounced effects on autonomic nervous system function or that cause sedation.

Despite earlier efforts to assess the variability in Irwin/FOB scoring and to identify influencing factors (14, 16, 24, 29), this is the first study to our knowledge investigating the effects of step-by-step harmonization on neuropharmacological safety assessment in laboratory mice. In this context, it should be noted that it is impossible to achieve a state of full harmonization in experimental preclinical animal studies on a multinational level. Full harmonization of animal housing conditions (e.g., day/night light cycle, temperature, humidity) can at best be within a predefined range. Moreover, a breeder with supply chains across continents can hardly be found. Therefore, the influence of genetic drift should be considered as a potential confounding factor in all stages of this study. Similarly, we could not ensure the full harmonization of drug manufacturers and batches due to specific manufacturer supply chains.

With respect to the harmonization of the scoring scheme, it is emphasized that the investigators for the newly introduced scoring scheme in stage 3 did not undergo a standardized training program. Different levels of training and professional experience of the investigators may have increased between-site variability. Due to the extensive study design, the capacities of the involved sites and different climate conditions across continents, a full harmonization of seasonal test conditions between sites could not be achieved. It is well known that behavioral patterns of laboratory rodents and *in-vivo* drug effects can be subject to considerable seasonal variation despite a standardized light/dark phase cycle under laboratory conditions (30, 31). Given that pharmacokinetics are also subject to seasonal influences, it is of interest that Himmel and colleagues (2019) identified drug exposure as a relevant source of variation in their Irwin/FOB multicenter study (24).

Considering the overall validity of this study as a multicenter approach, it needs to be considered that a relatively small number of participating laboratory sites (five sites for stages 1 and 2, respectively, and three sites for stage 3) was included.

Despite decades of efforts to develop standardized approaches for neurofunctional assessment (13, 25, 26, 32, 33), the current study confirmed relevant differences between variants of Irwin tests/FOBs used by different companies and institutes. In this context, it needs to be considered that Irwin/FOB data are an important element in safety pharmacology assessment (22). Reliable analysis of neurofunctional parameters is a presupposition for informed decision-making, prioritization, and de-risking during drug candidate selection processes before first-in-human testing (22, 34).

Considering the relevance of the test system, it is important to realize the high variability and poor reproducibility and to consider that the variants of the scoring systems and subjective scoring can have a major impact on the outcome. According to our findings, this applies not only to information about the extent, temporal course, and dose dependence of drug effects, but even to the mere detection of specific effects, i.e., to conclusions about their presence or absence. Pharmaceutical companies, contracting companies, and their clients need to be fully aware of the high variability and limited reproducibility.

The findings further underline the need for standardization, but it is also clear that scoring systems need to reflect the specificities of the indications and patient populations (e.g., pediatric, geriatric). In this regard, the parameters and scoring system should be carefully reviewed and adjusted if necessary.

Finally, our findings suggest that it can be useful to group different parameters into functional domains as a basis for comparison across laboratories with different scoring systems. The results of our meta-analysis demonstrated that grouping the parameters into the functional domains ‘motor’, ‘autonomic’, ‘sedation-based’, and ‘excitation-based’ reduced the heterogeneity of the dataset allowing comparative assessment.

Taken together, the findings of the multicenter study confirmed a pronounced interlaboratory variability in Irwin datasets for the CNS-active compound MK-801. Harmonization of various environmental and study design factors had no relevant effect on the qualitative and quantitative interlaboratory differences. In contrast, the study provided evidence that harmonization of the parameters and the scoring system can limit variability.

Companies, institutes, and experimenters need to carefully consider the high variability of Irwin/FOB data when making decisions during drug development.

## Methods

### Ethics statement

All animal experiments were conducted and reported in line with the EU Directive 2010/63/EU, the ARRIVE (Animal Research: Reporting of In Vivo Experiments) guidelines, and the Basel declaration (http://www.basel.declaration.org) including the 3R principle. Animal experiments were approved by the Government of Upper Bavaria (Munich, Germany, license number ROB-55.2-2532.Vet_02-18-45), the state investigation office Rheinland-Pfalz (Koblenz, Germany, license numbers 23 177-07/G14-9-082 and 23 177-07/G19-9-076), the Government Principle of Tübingen (license number 35/9185.81-8/18-025-G), the Project Authorization Board in the Regional State Administrative Agency for Southern Finland (license number 2018-22888), and the Institutional Animal Care and Use Committee (IACUC, license number 271_0315).

In addition, all sites adhered to the EQIPD critical principles for guiding the design, conduct, and analysis of preclinical efficacy and safety research.

### Experimental design

The experiment compared the variability of the Irwin test as a quick, commonly applied behavioral assay in preclinical safety pharmacology across five laboratory sites in Europe and the United States, including academic and industry sites. The five participating laboratory sites are listed below in alphabetical order. The identity of the laboratories has been masked to enable anonymized reporting of the results. Each laboratory was randomly assigned a number between 1 and 5. The number assigned to a laboratory was used for the entire analysis of the data from stages 1 to 3.

AbbVie (AbbVie Deutschland GmbH & Co KG, Ludwigshafen, Germany)

Boehringer Ingelheim (Boehringer Ingelheim Pharma GmbH & Co. KG, Biberach, Germany)

### LMU (Ludwig-Maximilians-Universität Munich, Munich, Germany)

Orion (Orion Corporation, Turku, Finland)

PsychoGenics (PsychoGenics Inc., New Jersey, USA)

The Irwin test was selected for this multicenter approach due to its frequent use in neuroscientific safety pharmacology. Besides potential side effects on the central nervous system, the Irwin score can also provide information on an animal’s general condition and the well-being of the laboratory animal. This multicenter approach comprised three stages with experimental protocols of a different, ascending degree of variable harmonization across sites.

### Experimental procedure

In each stage, the Irwin test was performed multiple times after acute compound administration to enable a comparison across treatment groups, dosages, and/or time. An overview of the experimental design is provided in **Figure 5**.

**Figure 5.**
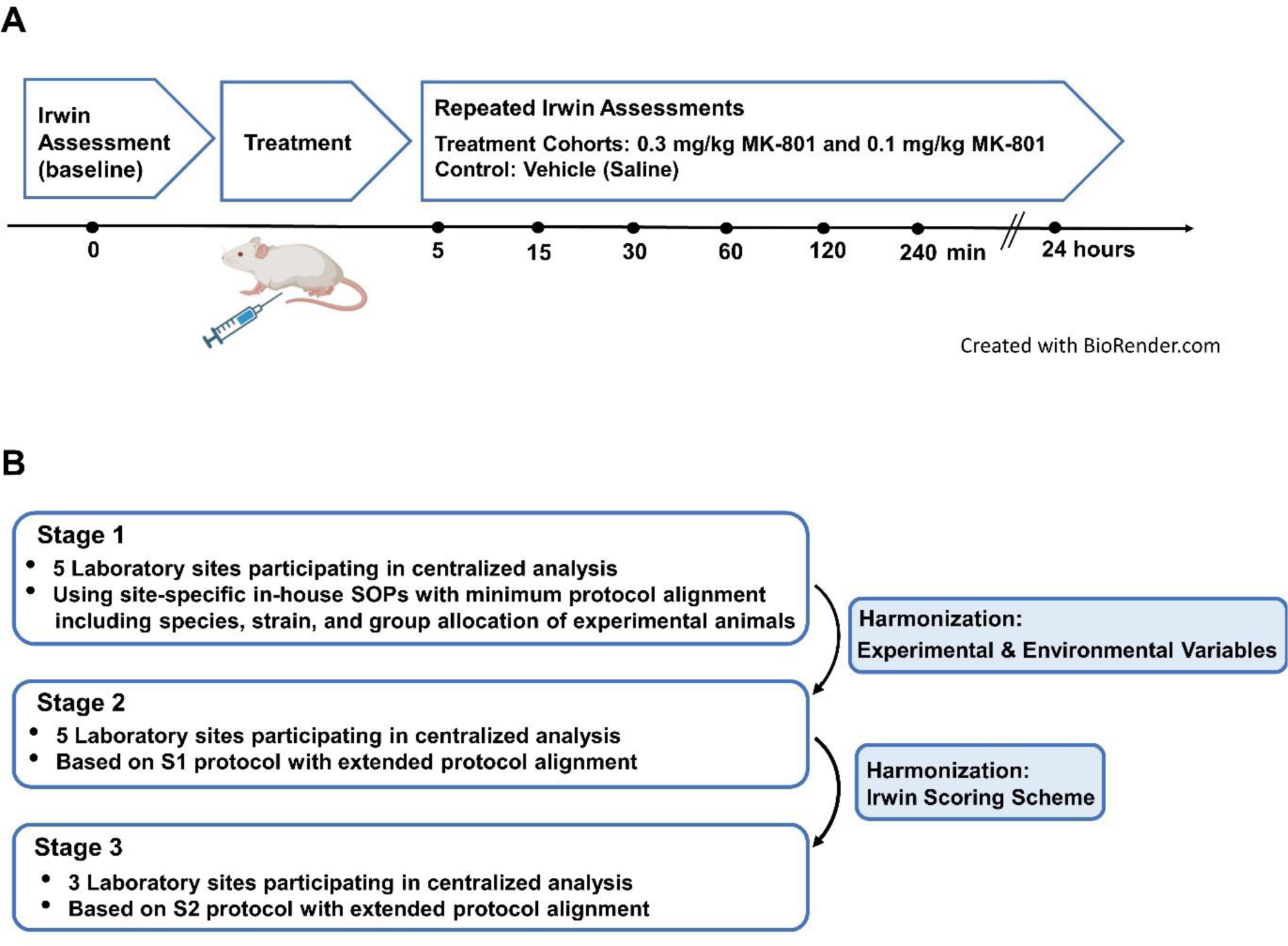
Overview of the experimental design and the stages of protocol harmonization across laboratory sites. **(A)** The local protocol used in stage 1 specified the following minimum requirements: species, strain, sex, age, treatment groups, and vehicle group. After an initial baseline assessment, Irwin tests were applied multiple times post-injection to assess the effects of MK-801, a non-competitive NMDA antagonist, over time. **(B)** In stage 2, experimental and environmental variables were harmonized across laboratory sites (**S12, S13, S14, S15, S16, S17, and S18 Supplementary Tables**) and the report of outcome measures was harmonized (**S19 Supplementary Table**) The experimental procedure in stage 3 was carried out using an Irwin scoring scheme shared across laboratory sites (**S20 Supplementary Table**).

Just before the experiments, the animals were housed individually. First, the Irwin test with each outcome measure was assessed at baseline. Immediately following baseline assessment, vehicle, 0.1 mg/kg MK-801, or 0.3 mg/kg MK-801 i.p. in a dosing volume of 10 ml/kg were administered. The compound and dosage did not vary between sites and stages. Details on the compounds used (trade name, manufacturer) are given in **S11 Supplementary Table**. A stopwatch was started at the time of injection. All outcome measures were assessed again 5, 15, 30, 60, 120, 240 minutes, and 24 hours post-injection. In deviation from the protocol, Irwin assessments 5 and 15 minutes after injections were omitted by site 2 in stage 1 and by site 3 in stage 2. When the 24-hour post-injection time point for all animals tested on the same day was completed, animals were returned to their original group housing cage.

The cage lid and mouse house were first removed for observations carried out in the home cage. For the handling observations, mice were tail-handled with gloved hands. During the wire maneuver, the mouse was held above a horizontal wire (diameter: 3.5 mm, 20 cm from table surface), lowered towards it to allow the forelimbs to grip and then held in extension and rotated around the wire. The placement of the forelimbs onto the wire was assessed. During visual placing, the mouse was lifted by the base of the tail to approximately 15 cm above a metal grid (1 cm grid size, raised 12.5 cm from table surface) and then lowered towards it in approximately 1-2 s, decelerating as the grid was approached. Extension of the forelimbs prior to reaching the grid was assessed. Observations in the general section were made throughout the duration of each testing time point. For the observations measured during restraint, the animals were briefly neck-restrained. The toe pinch reflex was assessed by pressing forceps between the digits of a hindlimb, and the corneal reflex was assessed by approaching the eye with a cotton swab. The righting reflex was assessed when releasing the mouse from restraint. To measure touch response, the mouse was gently stroked three times with the back of the experimenter’s hand. Startle response was assessed by loudly tapping a thick marker pen on the corner of the testing cage opposite from the mouse’s location.

### Protocol harmonization

During the first stage, also referred to as the ‘localization stage’, a minimum of requirements were set in terms of experimental and environmental factors. The local protocol specified the following minimum requirements: sex and strain (female and male NMRI mice), age (8-10 weeks), treatment (intraperitoneal (i.p.) injection of MK-801), dosage (0.1 mg/kg and 0.3 mg/kg), experimental groups (MK-801 0.1 mg/kg vs. MK-801 0.3 mg/kg), and vehicle group (saline). The Irwin test was applied multiple times after injection to assess the effects over time. The assessment timepoints were defined as follows: baseline assessment (just before the first injection), followed by assessments 5, 15, 30, 60, 120 minutes, and 24 hours post-injection. Apart from these predefined requirements given, each site performed the tests following their local in-house standard operating procedure (SOP). In this stage, five laboratory sites participated.

In the second stage, also referred to as the ‘harmonization stage’, a protocol was used to harmonize the parameters of the following main variables: 1) housing/husbandry (e.g. social housing, cage type, environmental enrichment, interaction with animals), 2) assignment to treatment groups including randomization, 3) personnel (i. e., experimenters supposed to be non-smokers, standardized handling method, blinding of experimenter to group allocation), and 4) experimental set-up (e.g., acclimatization, test arena, food/water restriction). Overviews of the main variables and parameters are provided in **S12, S13, S14 S15, S16, S17, and S18 Supplementary Tables**. A protocol about outcome measures nomenclatures and outcome measure rating systems was shared across sites to harmonize the report of scoring outcome measures (**S19 Supplementary Table**). In this stage, five laboratory sites participated.

In the third stage, also referred to as the ‘ring test stage’, a protocol was used that included complete harmonization of Iwrin scoring in addition to the variables harmonized during stage 2. An overview of the harmonized scoring scheme applied in stage 3 and their subsummation to functional domains is provided in **S20 Supplementary Table**. In this stage, four laboratory sites participated. As one laboratory site did not follow the shared protocol for a harmonized scoring scheme and did not collect baseline data, records from this lab were excluded from the centralized analysis. Details on the scoring scheme applied by lab 4 in stage 3 are provided in the **S21 and S22 Supplementary Table**.

### Experimental animals

Experiments were conducted with naïve mice during all stages. **Table 1** provides an overview of age, sex, housing, numbers, and dropouts of animals across sites and stages. Experimental animals were obtained from Charles River Laboratories (Sulzfeld, Germany), Taconic Biosciences, Inc. (Taconic’s Germantown, New York, USA), and Envigo RMS B.V. (Horst, the Netherlands). Further stage-specific information is provided in **S11 Supplementary Table**, including details on husbandry conditions (e.g., cage type, temperature, humidity, light cycle, frequency of cage change and cleaning, frequency of water/food replacement), experimental conditions (e.g., experimental set-up, analysis tools), and method of handling as well as number and gender of animal care takers. Details on blinding and randomization across sites and stages are provided in **S11 Supplementary Table**.

**Table 1.**
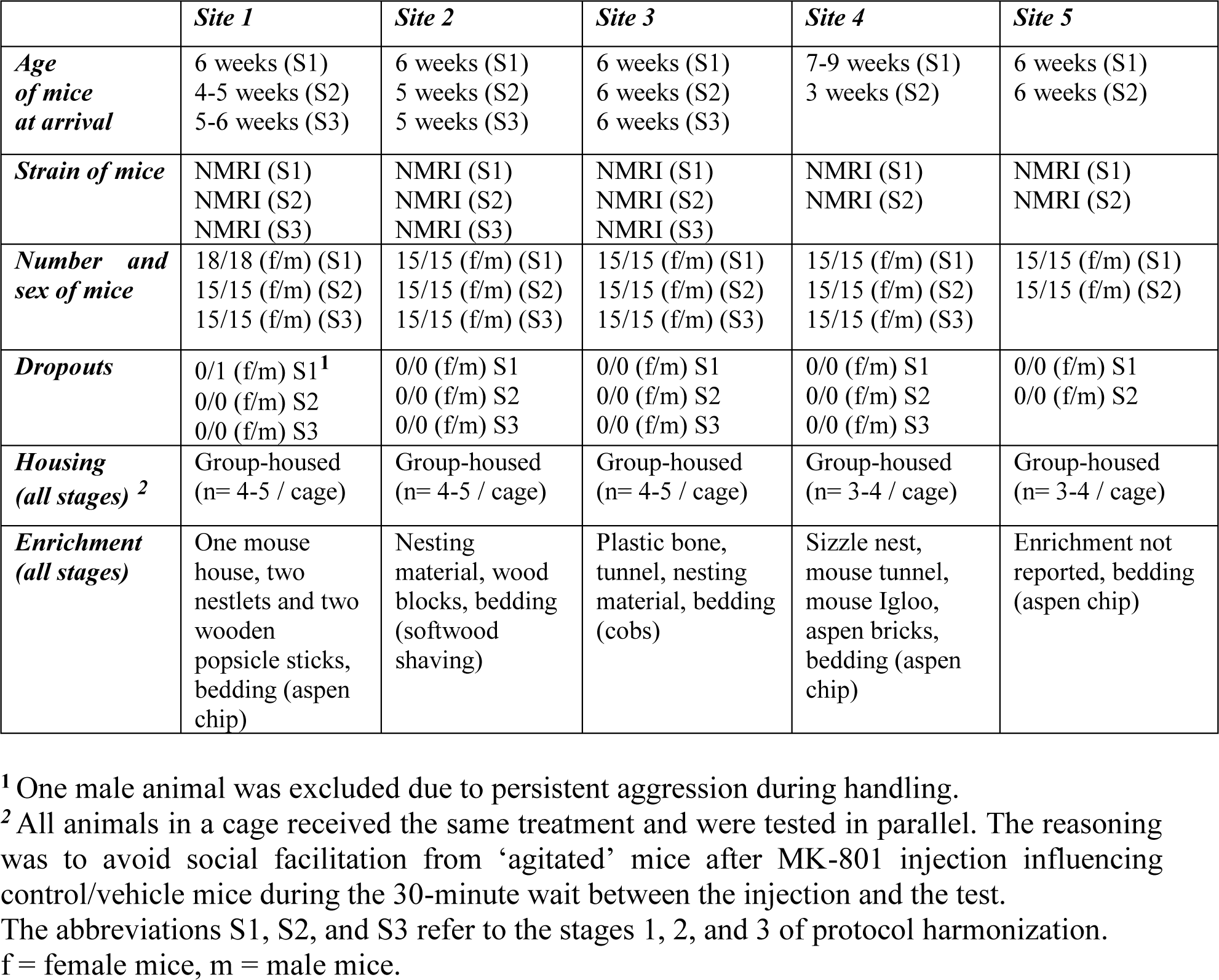
Details on experimental animals across the three stages of protocol harmonization.

|  | *Site 1* | *Site 2* | *Site 3* | *Site 4* | *Site 5* |
| --- | --- | --- | --- | --- | --- |
| ***Age of mice at arrival*** | 6 weeks (S1)<br>4-5 weeks (S2)<br>5-6 weeks (S3) | 6 weeks (S1)<br>5 weeks (S2)<br>5 weeks (S3) | 6 weeks (S1)<br>6 weeks (S2)<br>6 weeks (S3) | 7-9 weeks (S1)<br>3 weeks (S2) | 6 weeks (S1)<br>6 weeks (S2) |
| ***Strain of mice*** | NMRI (S1)<br>NMRI (S2)<br>NMRI (S3) | NMRI (S1)<br>NMRI (S2)<br>NMRI (S3) | NMRI (S1)<br>NMRI (S2)<br>NMRI (S3) | NMRI (S1)<br>NMRI (S2) | NMRI (S1)<br>NMRI (S2) |
| ***Number and sex of mice*** | 18/18 (f/m) (S1)<br>15/15 (f/m) (S2)<br>15/15 (f/m) (S3) | 15/15 (f/m) (S1)<br>15/15 (f/m) (S2)<br>15/15 (f/m) (S3) | 15/15 (f/m) (S1)<br>15/15 (f/m) (S2)<br>15/15 (f/m) (S3) | 15/15 (f/m) (S1)<br>15/15 (f/m) (S2)<br>15/15 (f/m) (S3) | 15/15 (f/m) (S1)<br>15/15 (f/m) (S2) |
| ***Dropouts*** | 0/1 (f/m) S1<sup>1</sup><br>0/0 (f/m) S2<br>0/0 (f/m) S3 | 0/0 (f/m) S1<br>0/0 (f/m) S2<br>0/0 (f/m) S3 | 0/0 (f/m) S1<br>0/0 (f/m) S2<br>0/0 (f/m) S3 | 0/0 (f/m) S1<br>0/0 (f/m) S2<br>0/0 (f/m) S3 | 0/0 (f/m) S1<br>0/0 (f/m) S2 |
| ***Housing (all stages) <sup>2</sup>*** | Group-housed<br>(n= 4-5 / cage) | Group-housed<br>(n= 4-5 / cage) | Group-housed<br>(n= 4-5 / cage) | Group-housed<br>(n= 3-4 / cage) | Group-housed<br>(n= 3-4 / cage) |
| ***Enrichment (all stages)*** | One mouse house, two nestlets and two wooden popsicle sticks, bedding (aspen chip) | Nesting material, wood blocks, bedding (softwood shaving) | Plastic bone, tunnel, nesting material, bedding (cobs) | Sizzle nest, mouse tunnel, mouse Igloo, aspen bricks, bedding (aspen chip) | Enrichment not reported, bedding (aspen chip) |

<sup>1</sup> One male animal was excluded due to persistent aggression during handling.
<sup>2</sup> All animals in a cage received the same treatment and were tested in parallel. The reasoning was to avoid social facilitation from ‘agitated’ mice after MK-801 injection influencing control/vehicle mice during the 30-minute wait between the injection and the test.
The abbreviations S1, S2, and S3 refer to the stages 1, 2, and 3 of protocol harmonization.
f = female mice, m = male mice.

### Data processing

Once data were acquired, each site transferred its raw data and metadata to an Excel file shared across sites, which was then used for centralized analysis.

First, scores from individual parameters were grouped into four domains, namely: 1) motor-functions related domain (‘motor domain’), 2) autonomic-system related domain (‘autonomic domain’), 3) excitation-related domain, and 4) sedation-related domain. Additional parameters were grouped as ‘other measures’, depending on the site-specific scoring schemes in stages 1 and 2. Second, change-from-baseline data were calculated to account for the individual condition of the animal and to mitigate the associated bias. The domain-specific Irwin sum scores assessed before injection (referred to as ‘baseline assessment’) were subtracted from the domain-specific sum scores measured at a time (5, 15, 30, 60, 120 minutes, and 24 hours post-injection, respectively). Third, the normalized sum scores were presented unidirectionally (positive values only) as delta sum scores.

Raw data and normalized data from each treatment group and vehicle group at each site across stages are provided in the Figshare Repository [DOI will be provided in the accepted article].

### Statistical analyses

The data processed as described above are illustrated per group, domain, time point of assessment, site, and stage as median with interquartile range (IQR), respectively.

A meta-analysis was conducted to compare effect sizes across sites within stage 3, using IBM SPSS Statistics (version 29.0). Change-from-baseline measures were recoded on a binary scale to allow for effect size calculation and meta-analysis of the scored data. For the binary classification of change scores, scores > 0 were given the value 1, while scores ≤ 0 were given the value 0. Effect sizes were calculated using the Odds Ratio (OR), based on the binary response of change scores of 0.1 mg/kg MK-801 vs. vehicle and 0.3 mg/kg MK-801 vs. vehicle. The meta-analysis was conducted using change scores 30 minutes post-injection, as the strongest effects of the injected compounds were measured at this time in all laboratories. The comparison of binary outcomes was based on the OR. Effect size averaging and weighing were based on a random effects model and the inverse variance method, including variance within and between studies. The variance was estimated using the REML method. In addition, subgroup analyses were conducted using the four functional domains as moderator variables. For each functional domain and outcome, Chi-Square test (Q statistics) assessed homogeneity in effect sizes to test whether the observed variability in effect sizes was unlikely to have arisen by chance. Standard error (SE) adjustment was not applied. We calculated standard deviations (SDs) when reporting p-values, t-values, SEs, and 95% CIs.

Analysis and graphical illustration of the processed data were performed with R, version 4.1.2, IBM SPSS Statistics for Windows, version 29.0, and GraphPad Prism for Windows, version 6.

## Supporting Information

**Appendix 1**

**Appendix 2**

## Funding

This project has received funding from the Innovative Medicines Initiative 2 Joint Undertaking under grant agreement No 777364. This Joint Undertaking receives support from the European Union’s Horizon 2020 research and innovation programme and the European Federation of Pharmaceutical Industries and Associations. Also, SRT was supported by the German Research Foundation (DFG) – research group FOR 2591 (BL953/11-1 and 11-2).

## Conflict of Interest Statement

The authors declare no conflicts of interest.

## Acknowledgments

We would like to thank Sarah Glisic (LMU), Katharina Gabriel (LMU), Sabine Vican (LMU), Uwe Roßberg (LMU), Anu Heino (Orion), Päivi Saikkonen (Orion), Michael Winter (Boehringer Ingelheim), Beatrice Kley (Boehringer Ingelheim), and Juliane Denz (Boehringer Ingelheim) for their contributions to the project.

This publication reflects only the authors’ view and the Innovative Medicines Initiative 2 Joint Undertaking is not responsible for any use that may be made of the information it contains.

## Data Availability Statement

The data used in the current study are publicly available in the Figshare Repository [DOI will be provided in the accepted article]. All other summaries and raw data are within the paper and its Supplementary files.

## Author Contributions

Maria Reiber: data processing, formal analysis, review and editing of the draft, visualization, writing of the original draft

Helen Stirling: data processing, investigation, review and editing of the draft, visualization

Tim P. Ahuis: data processing, investigation

Washington Arias: investigation

Katharina Aulehner: investigation, methodology, study design

Ute Dreßler: investigation

Martien J.H. Kas: conceptualization, funding acquisition, methodology, review and editing of the draft, study design

Johanna Kela: supervision

Kimberly Kerker: data processing

Tarja Kuosmanen: investigation

Helga Lorenz: data processing, supervision

Alexander T. Pennington: investigation

Eva-Lotta von Rüden: study design, supervision

Heike Schauerte: methodology, supervision

Isabel Seiffert: investigation

Steven R. Talbot: review and editing of the draft, supervision

Christina Torturo: supervision

Sami Virtanen: data processing

Ann-Marie Waldron: investigation

Sylvie Ramboz: conceptualization, data processing, funding acquisition, methodology, project administration, review and editing of the draft, study design, supervision

Heidrun Potschka: conceptualization, funding acquisition, methodology, review and editing of the draft, study design, supervision, visualization, writing of the original draft

## Notes

### Competing Interest Statement

The authors have declared no competing interest.

