## Appendix 1 for "A Systematic Assessment of Robustness in CNS Safety Pharmacology"

### *Appendix 1: Supplementary Figures*

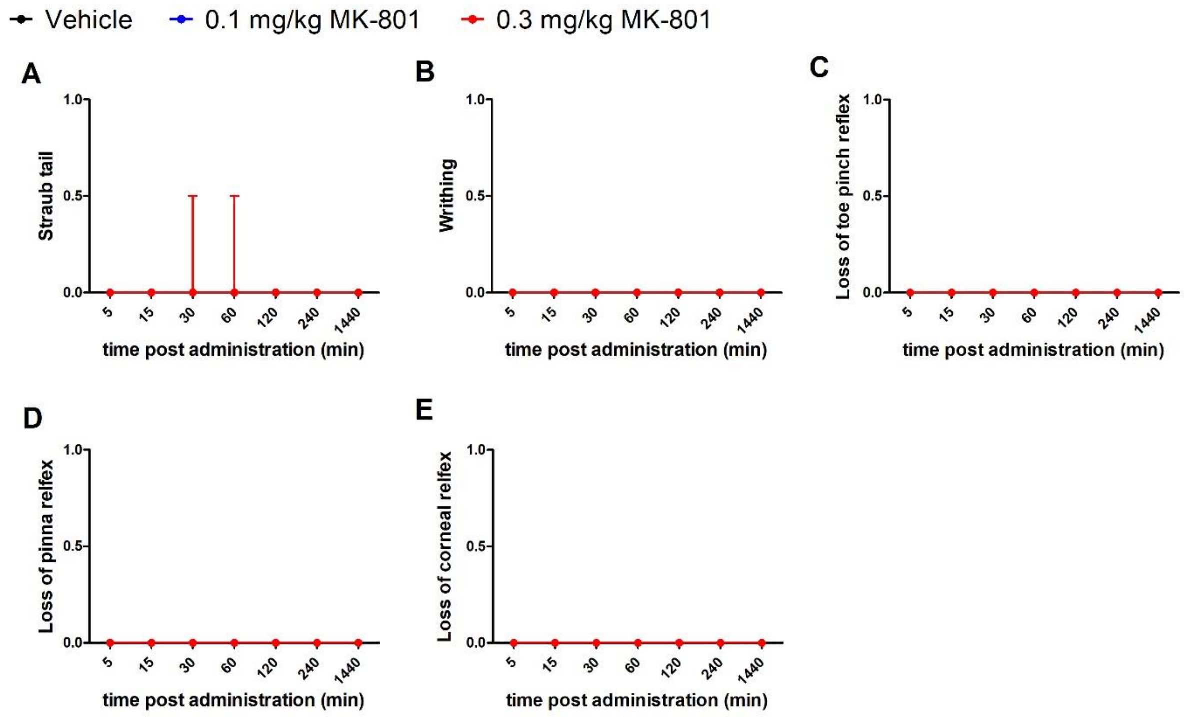


**S1 Figure. Visualization of stage 1 data 'other measures' as provided by site 1.**
Data are presented as delta sum scores per timepoint (positive change-from-baseline scores). Data are presented as median with interquartile range (IQR). Details about the site-specific scoring scheme are provided in **S1 and S2 Supplementary Tables**.

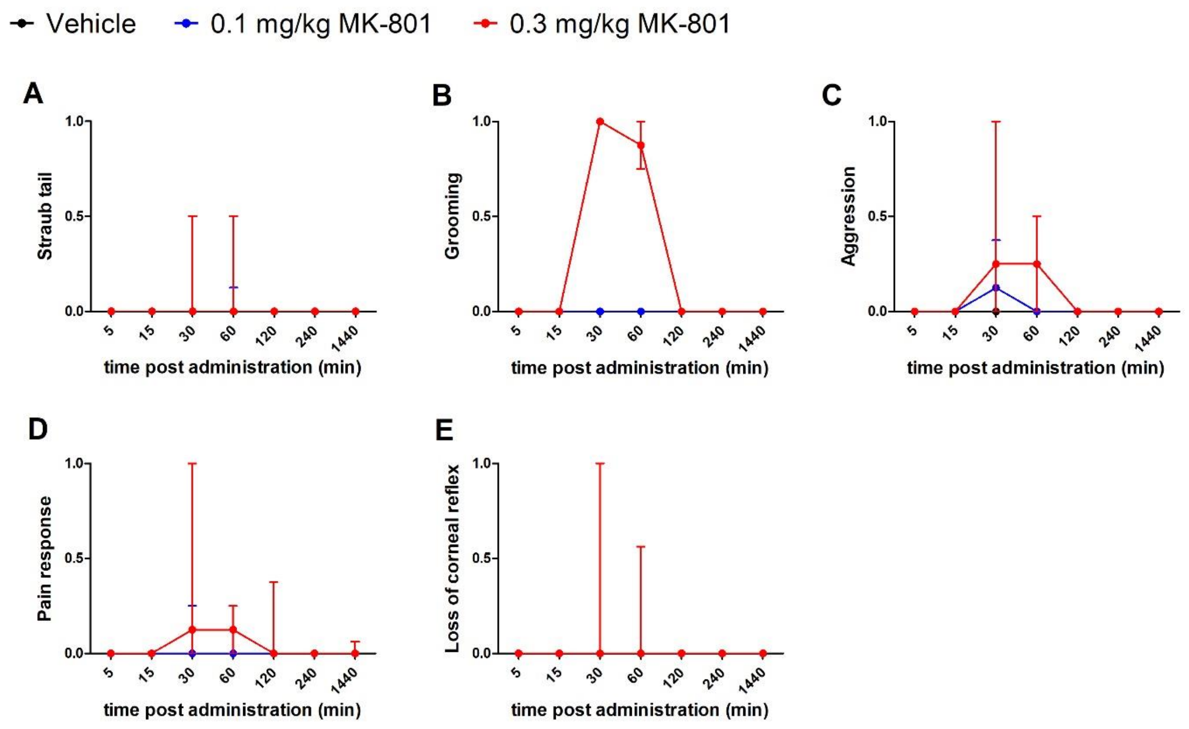


**S2 Figure. Visualization of stage 1 data ‘other measures’ as provided by site 2.**
Data are presented as delta sum scores per timepoint (positive change-from-baseline scores). Data are presented as median with interquartile range (IQR). Details about the site-specific scoring scheme are provided in **S3 and S4 Supplementary Tables**.

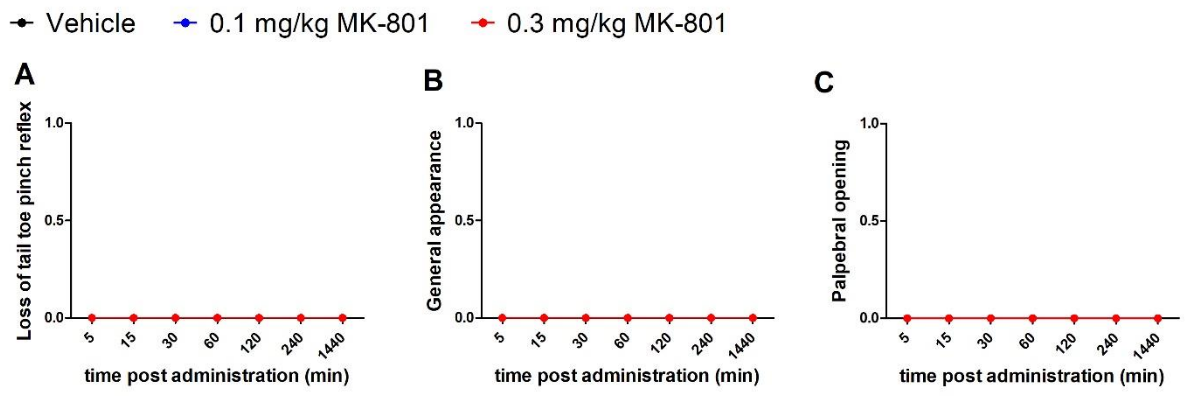


**S3 Figure. Visualization of stage 1 data ‘other measures’ as provided by site 3.**
Data are presented as delta sum scores per timepoint (positive change-from-baseline scores). Data are presented as median with interquartile range (IQR). Details about the site-specific scoring scheme are provided in **S5 and S6 Supplementary Tables**.

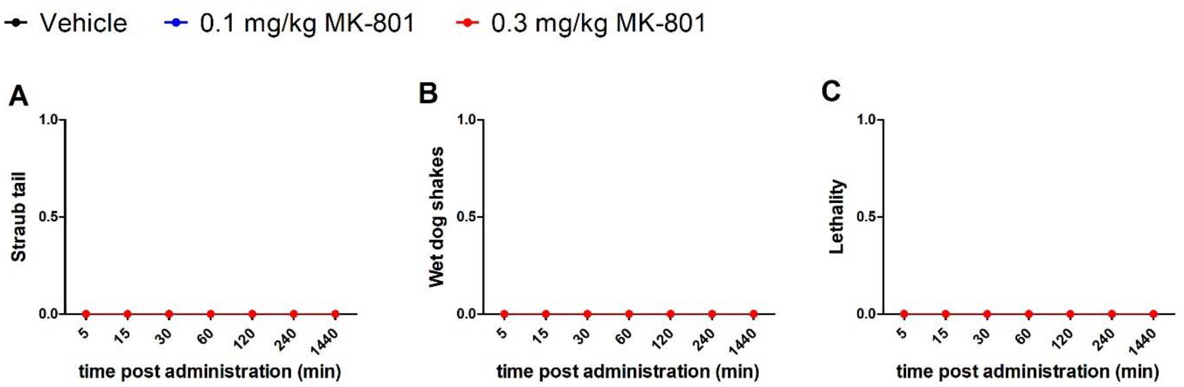


**S4 Figure. Visualization of stage 1 data 'other measures' as provided by site 4.**
Data are presented as delta sum scores per timepoint (positive change-from-baseline scores). Data are presented as median with interquartile range (IQR). Details about the site-specific scoring scheme are provided in **S7 and S8 Supplementary Tables**.

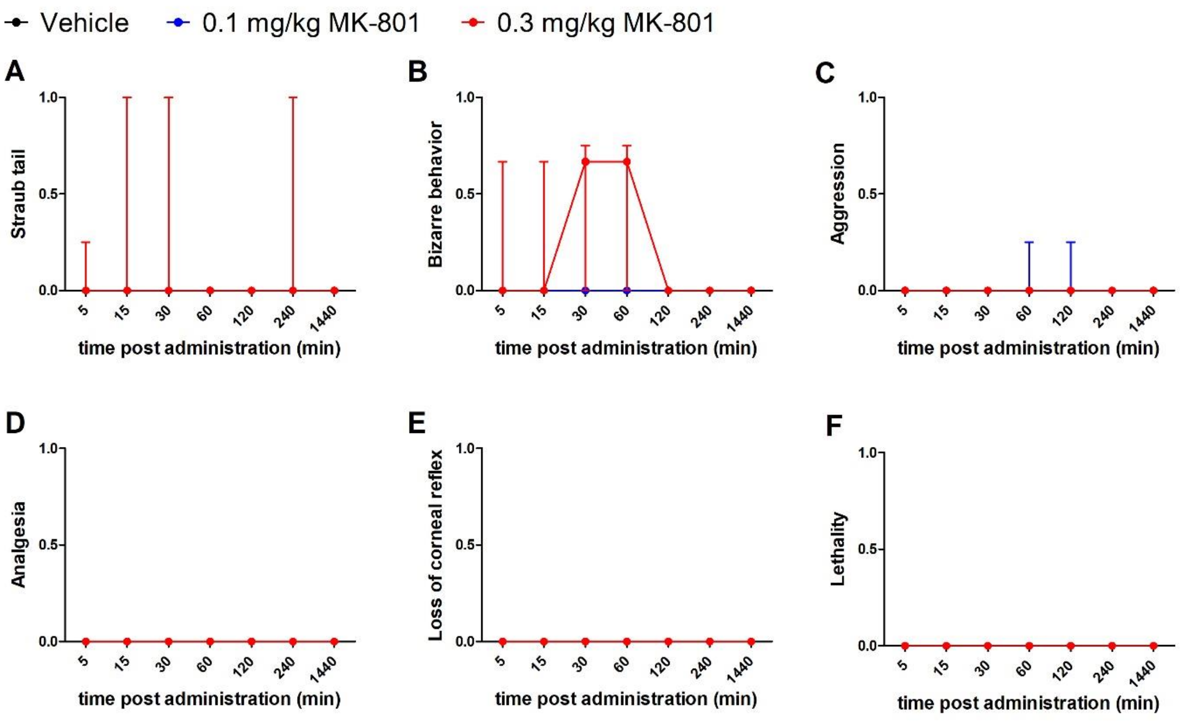


**S5 Figure. Visualization of stage 1 data 'other measures' as provided by site 5.**
Data are presented as delta sum scores per timepoint (positive change-from-baseline scores). Data are presented as median with interquartile range (IQR). Details about the site-specific scoring scheme are provided in **S9 and S10 Supplementary Tables**.

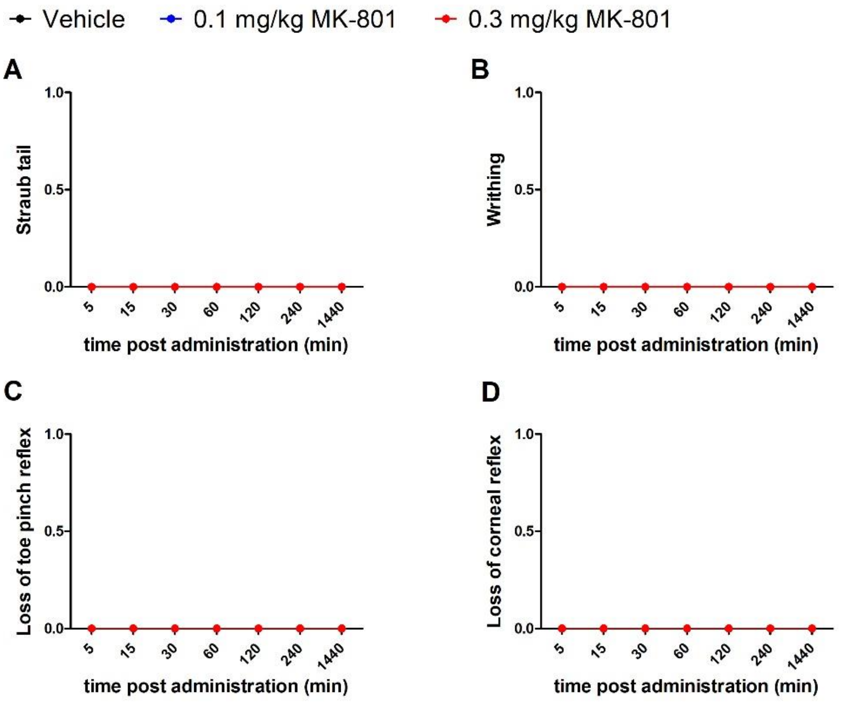


**S6 Figure. Visualization of stage 2 data 'other measures' as provided by site 1.**
Data are presented as delta sum scores per timepoint (positive change-from-baseline scores). Data are presented as median with interquartile range (IQR). Details about the site-specific scoring scheme are provided in **S19 Supplementary Table**.

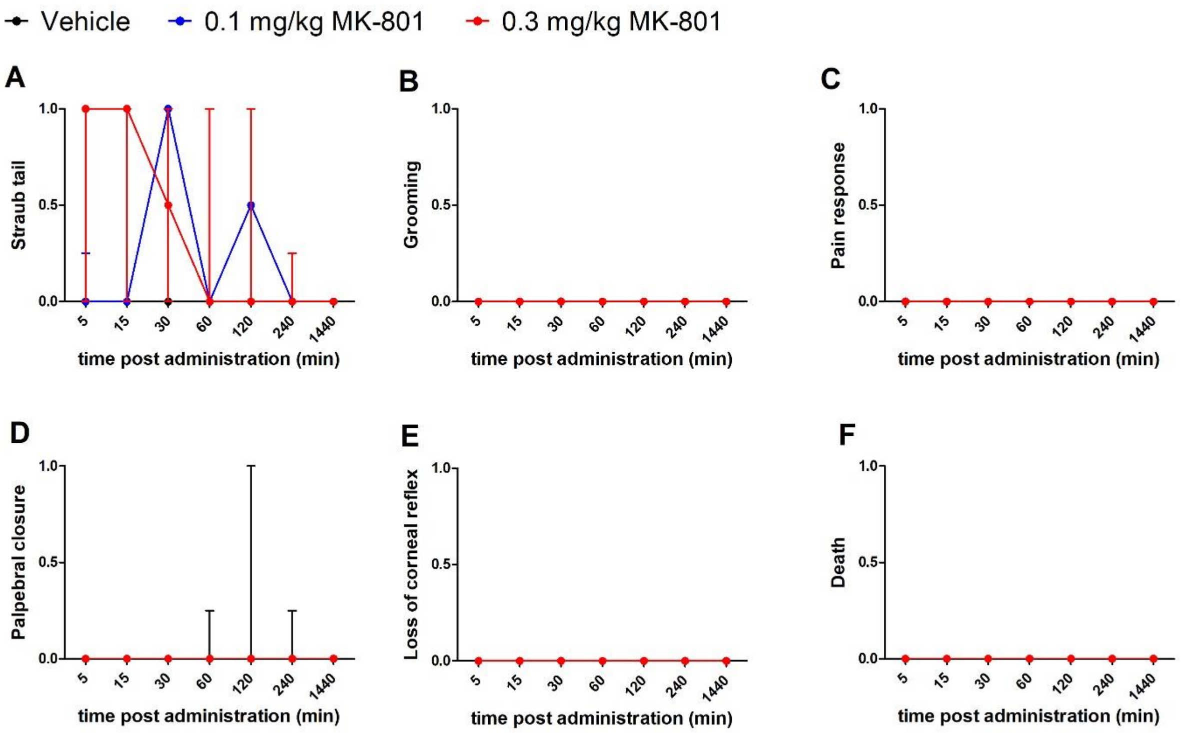


**S7 Figure. Visualization of stage 2 data ‘other measures’ as provided by site 2.**
Data are presented as delta sum scores per timepoint (positive change-from-baseline scores). Data are presented as median with interquartile range (IQR). Details about the site-specific scoring scheme are provided in **S19 Supplementary Table**.

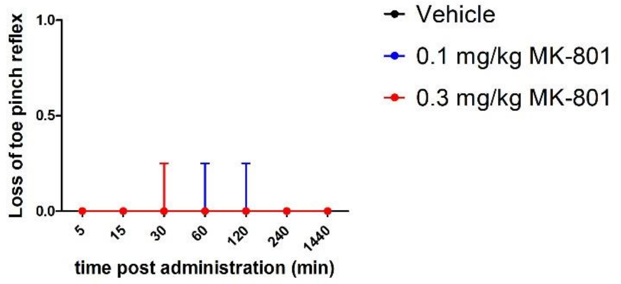


**S8 Figure. Visualization of stage 2 data ‘other measures’ as provided by site 3.**
Data are presented as delta sum scores per timepoint (positive change-from-baseline scores). Data are presented as median with interquartile range (IQR). Details about the site-specific scoring scheme are provided in **S19 Supplementary Table**.

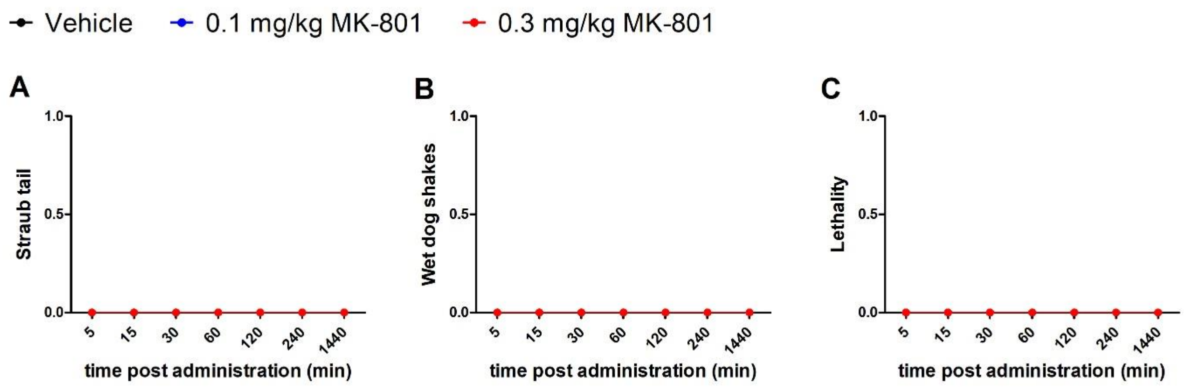


**S9 Figure. Visualization of stage 2 data 'other measures' as provided by site 4.**
Data are presented as delta sum scores per timepoint (positive change-from-baseline scores). Data are presented as median with interquartile range (IQR). Details about the site-specific scoring scheme are provided in **S19 Supplementary Table**.

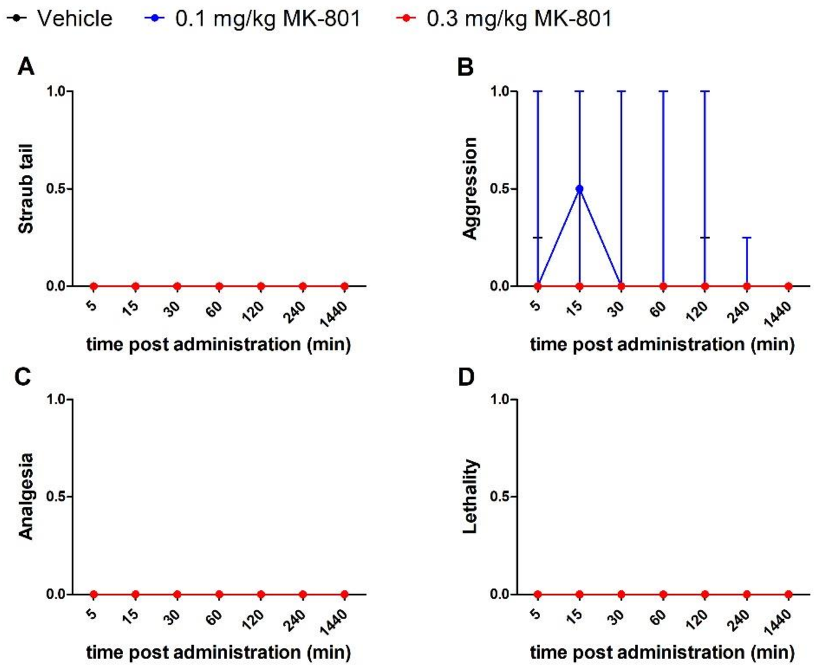


**S10 Figure. Visualization of stage 2 data 'other measures' as provided by site 5.**
Data are presented as delta sum scores per timepoint (positive change-from-baseline scores). Data are presented as median with interquartile range (IQR). Details about the site-specific scoring scheme are provided in **S19 Supplementary Table**.

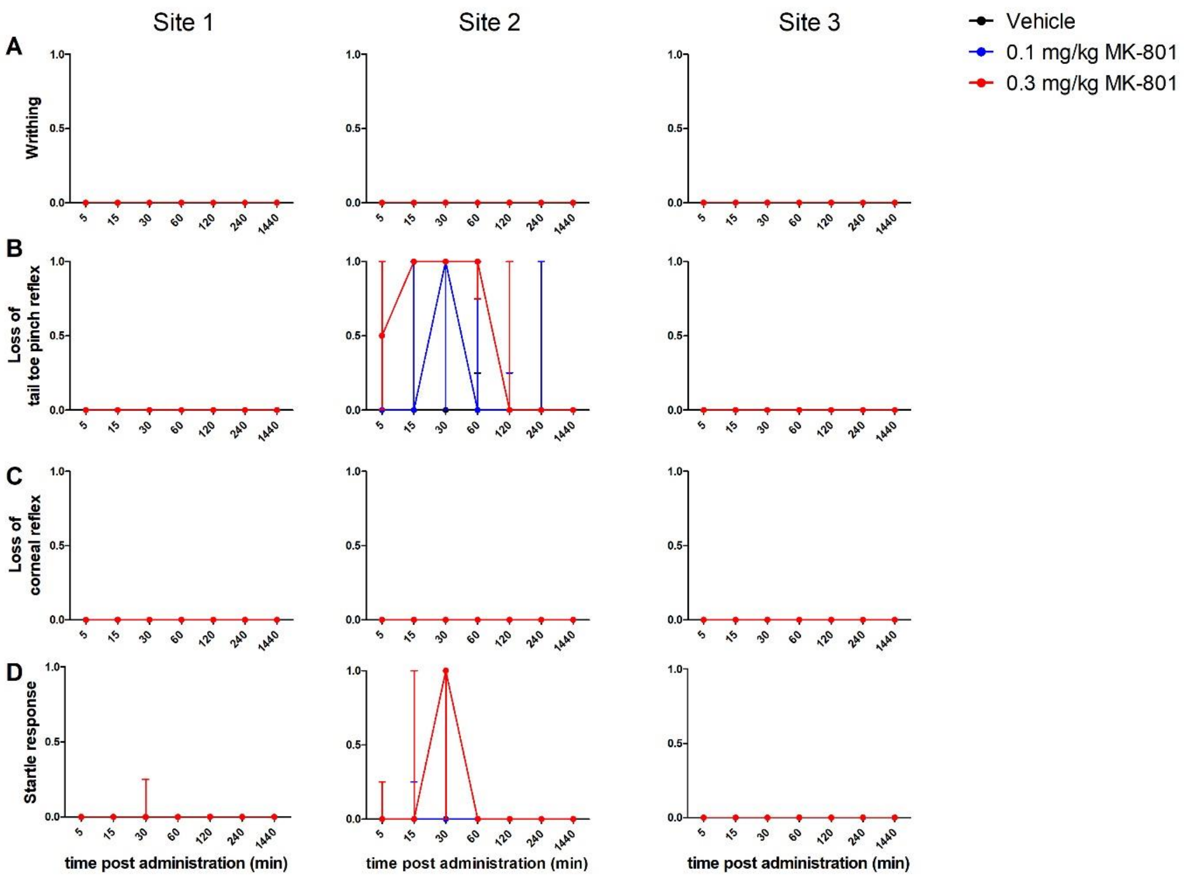


**S11 Figure. Visualization of stage 3 data 'other measures' as provided by site 1, 2, and 3.** In accordance with the shared stage 3 protocol, the following harmonized outcome measures are reported: A) Writhing, B) Loss of tail/toe pinch reflex, C) Loss of corneal reflex, and D) Altered startle response. Data are presented as delta sum scores per timepoint (positive change-from-baseline scores). Data are presented as median with interquartile range (IQR). Details about the shared protocol are provided in **S20 Supplementary Table**.

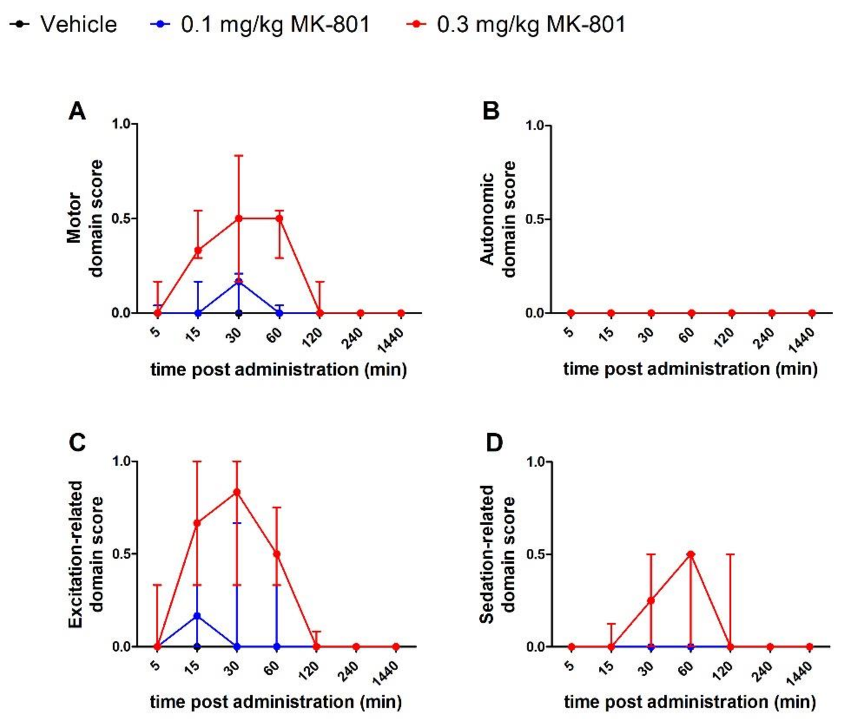


**S12 Figure. Visualization of stage 3 data as provided by site 4.**
Data are presented as sum scores per timepoint for each of the four functional domains. The stage 3 data provided by site 4 do not include baseline measurements. Therefore, the present data cannot be presented as delta sum scores per timepoint (positive change-from-baseline scores). Data are presented as median with interquartile range (IQR). Details about the scoring scheme applied by site 4 are provided in **S21 Supplementary Table**.

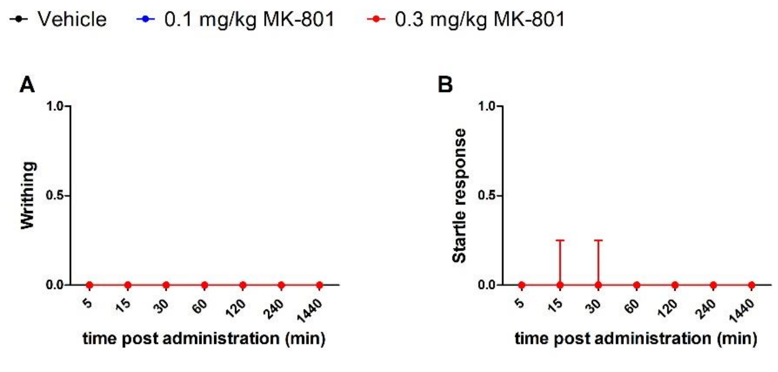


**S13 Figure. Visualization of stage 3 data ‘other measures’ as provided by site 4.**
Data are presented as sum scores per timepoint. Stage 3 data as provided by site 4 do not include baseline measurements. Therefore, the present data cannot be presented as delta sum scores per timepoint (positive change-from-baseline scores). Data are presented as median with interquartile range (IQR). Details about the scoring scheme applied by site 4 are provided in **S21 Supplementary Table**.
