## Appendix 2 for "A Systematic Assessment of Robustness in CNS Safety Pharmacology"

### *Appendix 2: Supplementary Tables*

**S1 Table. Overview of the experimental stage 1 scoring parameters as provided by site 1 and their allocation to the four functional domains for data analysis across sites.**
The results of the parameters allocated to the four functional domains are presented in **Figure 1A, B, C, D**. The results of the parameters subsumed under 'other measures' are presented in **S1 Supplementary Figure**.

| *Parameter* | *Scoring Range* |
|---|---|
| *Motor Domain* | |
| Grip strength | [-2]-[+1] |
| Wire manouvre | [0]-[-2] |
| Visual placing | [0]-[+2] |
| Abnormal gait | [0]-[+2] |
| Limb position | [0]-[1] |
| Righting reflex | [0]-[-2] |
| Pelvic elevation | [-1]-[+2] |
| Body tone | [-2]-[+2] |
| Limb tone | [-1]-[+1] |
| *Autonomic Domain* | |
| Piloerection | [0]-[1] |
| Exophthalmoses | [0]-[1] |
| Lacrimation | [0]-[1] |
| Salivation | [0]-[1] |
| Urination | [0]-[1] |
| Defecation | [0]-[1] |
| Body temperature | [-1]-[+1] |
| Skin color | [-2]-[+2] |
| Respiration | [-1]-[+1] |
| *Excitation-related Domain* | |
| Stereotypy | [0]-[1] |
| Tremors | [0]-[1] |
| Twitches | [0]-[1] |
| Convulsions | [0]-[1] |
| Vocalization | [0]-[1] |
| Irritability | [0]-[1] |
| Alertness | [-2]-[+2] |
| Spontaneous activity | [-1]-[+2] |
| Startle response | [-1]-[1] |
| Touch response | [-2]-[+2] |
| Fearfulness | [-1]-[+1] |
| *Sedation-related Domain* | |
| Passivity | [0]-[-2] |
| Palpebral opening | [-1]-[+1] |
| Alertness | [-2]-[+2] |
| Spontaneous activity | [-2]-[+2] |

| | |
|---|---|
| Startle response | [-1]-[1] |
| Touch response | [-2]-[+2] |
| Fearfulness | [-1]-[+1] |
| *Other Measures* | *Scoring Range* |
| Straub tail | [0]-[3] |
| Toe pinch reflex | [0]-[-2] |
| Pinna reflex | [0]-[-2] |
| Corneal reflex | [0]-[-2] |
| Writhing | [0]-[1] |

**S2 Table. Details about the stage 1 scoring scheme as provided by site 1.**

| *Outcome Measures* | *Scoring Range* | *Description* |
|---|---|---|
| Piloerection | [0]-[1] | Raised fur |
| Alertness | [-2]-[+2] | Relates to excitation/sedation |
| Writhing | [0]-[1] | Visually apparent abdominal movements e.g. stretching hind limbs with hollow flanks |
| Stereotypy | [0]-[1] | Absent or present, type of sterotypy given separately |
| Straub tail | [0]-[3] | Condition in which an animal carries its tail in an erect (vertical or nearly vertical) position |
| Tremors | [0]-[1] | Absent or present |
| Twitches | [0]-[1] | Absent or present |
| Convulsions | [0]-[1] | Absent or present, type of convulsion given separately |
| Fearfulness | [-1]-[+1] | Fear in response to the investigator's hand slowly approaching |
| Vocalization | [0]-[1] | In response to handling |
| Irritability | [0]-[1] | Propensity to bite when handled |
| Passivity | [0]-[-2] | Normal level of struggle, diminshed or absent |
| Grip strength | [-2]-[+1] | Diminished, normal or increased grip strength |
| Body tone | [-2]-[+2] | Estimated by noting the muscle tension by palpation. From Flaccid to rigid. |
| Toe pinch reflex | [0]-[-2] | Normal, diminshed or absent |
| Corneal reflex | [0]-[-2] | Approach eye with cotton swab - does it close eyes? |
| Wire maneouver | [0]-[-2] | The mouse should place both hindlimbs on the wire when rotated. Normal, diminished or absent. |
| Visual Placing | [0]-[+2] | Reflex is intact, or extent to which it is diminished |
| Pupil size | [-1]-[+1] | Constricted, normal, dilated |
| Palpebral opening | [-1]-[+1] | Narrowed, normal or widened |
| Exophthalmoses | [0]-[1] | Eyeball protrusion |
| Lacrimation | [0]-[1] | Visible dampness around mouth |
| Salivation | [0]-[1] | Visible dampness around eyes |
| Urination | [0]-[1] | Urination in response to handling |
| Defecation | [0]-[1] | Defecation in response to handling |
| Body temperature | [-1]-[+1] | Increased or decreased |
| Respiration | [-1]-[+1] | Increased or decreased |
| Skin color | [-2]-[+2] | From blanching to deep red |
| Limb tone | [-1]-[+1] | Grasping a forepaw of the mouse and noting the resistance to the extension of the paw. Normal, decreased or increased resistance |
| Pinna reflex | [0]-[-2] | |
| Righting reflex | [0]-[-2] | Place animal on back - see how it rights itself. Diminished, normal or completely absent |
| Spontaneous activity | [-2]-[+2] | Hypo- or hyper- locomotion |
| Pelvic elevation | [-1]-[+2] | From flat to highly elevated |
| Limb position | [0]-[1] | Abnormal rotation of limb |
| Abnormal gait | [0]-[+2] | Motor incoordination, ataxia. |
| Touch response | [-2]-[+2] | Flight reaction to repeated stroking on back with finger. Diminshed, normal or exaggerated response |
| Startle response | [-1]-[1] | Response to the noise of a pen tapped off the side of the cage. Diminshed, normal or exaggerated response |

**S3 Table. Overview of the experimental stage 1 scoring parameters as provided by site 2 and their allocation to the four functional domains for data analysis across sites.**

The results of the parameters allocated to the four functional domains are presented in **Figure 1A, B, C, D**. The results of the parameters subsumed under 'other measures' are presented in **S2 Supplementary Figure**.

| *Parameter* | *Scoring Range* |
|---|---|
| *Motor Domain* | |
| Gait | [0-8] |
| Grip strength | [0-8] |
| Limb position | [0-8] |
| Body posture | [0-8] |
| Catalepsy | [0-4] |
| Extensor reflex | [0-8] |
| Flexor reflex | [0-8] |
| *Autonomic Domain* | |
| Piloerection | [0-4] |
| Lacrimation | [0-4] |
| Skin colour | [0-4] |
| Salivation | [0-4] |
| Pupil size | [0-8] |
| Respiration | [0-8] |
| Diarrhoea | [0-4] |
| En-/Exophthalmus | [0-8] |
| *Excitation-related Domain* | |
| Stereotypies | [0-1] |
| Tremor | [0-4] |
| Vocalization | [0-4] |
| Convulsions | [0-4] |
| Spontaneous activity | [0-8] |
| Irritability | [0-8] |
| Struggle response | [0-8] |
| Induced activity | [0-8] |
| *Sedation-related Domain* | |
| Palpebral closure | [0-8] |
| Spontaneous activity | [0-8] |
| Irritability | [0-8] |
| Struggle response | [0-8] |
| Induced activity | [0-8] |
| *Other Measures* | |
| Straub tail | [0-4] |
| Corneal reflex | [0-8] |
| Grooming | [0-8] |
| Aggression | [0-4] |
| Pain response | [0-8] |

**S4 Table. Details about the stage 1 scoring scheme as provided by site 2.**

| *Outcome Measure* | *Scoring Range* | *Description* |
|---|---|---|
| Aggression | [0-4] | Aggression (normal = 0)<br>The intensity is scored from 0 to 4 |
| Body posture | [0-8] | Body posture (normal = 4)<br>0 – Is able to lie on its back (Animal is lying on its side or belly and one is able to move it to lie on its back)<br>1 – Lying on its side<br>2 – Lying on its belly<br>3 – Crouched together<br>4 – Normal sitting position<br>5 – Rearing<br>6- Walking around<br>7 – Restless<br>8 – "Frolicking" – jumping and running around |
| Catalepsy | [0-4] | Catalepsy (normal = 0)<br>0. No catalepsy<br>1 – 3. Cataleptic in increasing intensity<br>4. Animal is completely cataleptic – manipulation in different positions and postures is possible<br>Flaccid forms of catalepsy can be characteristic of a certain substance, if so one can add an S to the score |
| Convulsions | [0-4] | Convulsions (normal = 0)<br>1. Lifting front extremities and occasional twitching<br>2. Multiple spasms of hind extremities<br>3. Generalized seizure<br>4. Repeated generalized seizures of both fore and hind limbs<br>The predominant type of convulsion to be classified as K for clonic and T for tonic. |
| Corneal reflex | [0-8] | Corneal reflex (normal =4)<br>The strength of the reaction is scaled from 0 – 8<br>0. No reaction<br>1. Delayed/slowed down reaction<br>4. Normal reaction<br>8. Heightened reaction |
| Diarrhoea | [0-4] | Diarrhea (normal = 0)<br>Amount of water content is scored from 0 – 3 and 4 indicating pulpy feces |
| En-/Exophthalmus | [0-8] | En-/Exophthalmos (Normal = 4)<br>0 – 3. Enophthalmos in increasing intensity<br>4 – Normal position of the eyeball in the orbital bone<br>5 – 8. Protrusion of eyeball from the orbital bone in increasing intensity – with 8 meaning bulging eyes |
| Extensor reflex | [0-8] | Extensor reflex (Normal = 4)<br>Intensity scored 0 - 8 |
| Flexor reflex | [0-8] | Flexor reflex (Normal = 4)<br>Intensity scored 0 – 8 |

| *Outcome Measure* | *Scoring range* | *Description* |
|---|---|---|
| Gait | [0-8] | Gait (normal = 4)<br>0. Paralysis<br>1. Prone position<br>2. Unstable, staggering motion<br>3. Weak/Slack<br>4. Normal gait<br>5. Stretched/high legged walking<br>6. Toe walking<br>7 - 8. Spastic paralysis with splayed limbs |
| Grip strength | [0-8] | Grip strength (Normal = 4)<br>Intensity scored 0-4 (Tested by ability to hold on to inverted cage lid) |
| Grooming | [0-8] | Grooming (normal = 4)<br>0 – 3 Reduced grooming behavior, 4 Normal grooming<br>5 – 8 Increased grooming behavior |
| Induced activity | [0-8] | Spontaneous (a) and induced (b) activity (normal = 4)<br>0. Normal/resting position – no movement<br>1. Horizontal sniffing movements at rest<br>2. Lifting head at rest<br>3. Walking slowly<br>4. Normal behavior with grooming, head lifting and walking around<br>5. Frequently running around<br>6. Standing up straight, rearing and raising tail<br>7. Restlessness and running around a lot<br>8. Jumping around and whipping tail |
| Irritability | [0-8] | Irritability and flight (normal = 4)<br>0. No reaction<br>1. Closing eyes and cower away<br>2. Eyelid closing and slow evasion<br>3. Evasive movements<br>4. Normal reaction – retreat, moving ears towards body and grooming head<br>5. Standing upright and intensive head grooming<br>6. Head shaking and strong evasive movements<br>7. Violently jumping away and vocalizations<br>8. Jump up followed by running around and strong vocalizations |
| Lacrimation | [0-4] | Lacrimation (normal = 0)<br>Scored from 0 – 4 with 4 denoting a particularly strong secretion of fluid. |
| Limb position | [0-8] | Limb position (Normal = 4)<br>0 – 3 Extremities, especially the hind ones are stretched to the side or back, mostly limp<br>4 Extremities are in normal position under the body<br>5 – 8 Extremities are retracted and tense |

| *Outcome Measures* | *Scoring range* | *Description* |
|---|---|---|
| Pain response | [0-8] | Pain response (Normal = 4)<br>0. No pain reaction<br>1. Weak vocalization<br>2. Flight movements<br>3. Strong vocalization & flight movements<br>4. Normal pain reaction<br>5. Vigorously attacking<br>6. Running around<br>7. Biting itself and littermates<br>8. Wildly attacking and jumping up high |
| Palpebral closure | [0-8] | Paplebral closure (normal = 4)<br>0 – 3. Small gap, if extreme eyelids are tightly pinched together<br>4. Normal width compared to untreated animals<br>5 – 8. Eyes wide open giving the appearance that the eyes are swelled up |
| Piloerection | [0-4] | Piloerection (normal = 0)<br>scored from 0 – 4. |
| Pinna reflex | [0-8] | Pinna (ear muscle) reflex (normal = 4)<br>Intensity scored 0 - 8 |
| Pupil size | [0-8] | Pupil size (normal = 4)<br>The pupil size is assessed under a binocular lamp with small magnification.<br>0 – 3. Pupil shrinking<br>4. Normal pupil size<br>5 -8. Dilation of pupil |
| Respiration | [0-8] | Respiration (normal = 4)<br>0. Breathlessness<br>1 – 3. Slowing of breathing graded according to intensity<br>4. Normal breathing<br>5 – 8. Accelerated breathing |
| Salivation | [0-4] | Salivation/Drooling (normal = 0)<br>Scored from 0 – 4 with 4 denoting very strong salivation. |
| Skin colour | [0-4] | Skin color (normal = 0)<br>0. Normal color<br>1. Pale white<br>2. Dark red<br>3. Pale cyanotic<br>4. Dark cyanotic |

| *Outcome Measures* | *Scoring Range* | *Description* |
|---|---|---|
| Spontaneous activity | [0-8] | Spontaneous (a) and induced (b) activity (normal = 4)<br>0. Normal/resting position – no movement<br>1. Horizontal sniffing movements at rest<br>2. Lifting head at rest<br>3. Walking slowly<br>4. Normal behavior with grooming, head lifting and walking around<br>5. Frequently running around<br>6. Standing up straight, rearing and raising tail<br>7. Restlessness and running around a lot<br>8. Jumping around and whipping tail |
| Stereotypies | [0-1] | Stereotypies (normal = 0)<br>The intensity of stereotypic behaviors is scored from 0 – 4, the following types can be distinguished and noted<br>a. Grooming/ cleaning head (P)<br>b. Licking (L)<br>c. Jaw movements (Ki)<br>d. Paw movements (Pf)<br>e. Scratching (Kr)<br>f. Shaking head (S)<br>g. Turning head (D)<br>h. Tail whipping (SS)<br>i. Running around in circles (M)<br>j. Biting itself (Sb)<br>k. Picking up shavings/wood chippings (AS)<br>l. Jumping movements (Sp)<br>m. Sniffing movements (Sch)<br>Other stereotypies such as "rock and roll", walking backwards and the like must be noted in an extra box |
| Straub tail | [0-4] | Straub Tail (Normal = 0)<br>intensity scored from 0 – 4. |
| Struggle response | [0-8] | Struggle response/defense behavior (Normal = 4)<br>0. Raising the mouse on a hind paw is possible, tail is hanging down limply<br>1. Raising the mouse on a front limb is possible, tail is hanging down limply<br>2. Animal lies calmly on the hand<br>3. Lifting by the neck skin lightly is possible<br>4. Normal reaction to being scruffed<br>5 – 6. Animal is difficult to scruff, only with extra force, flees<br>7 – 8. Animal struggles, throws itself on its back or attacks the hand |
| Tremor | [0-4] | Tremor (Normal = 0)<br>intensity is scored from 0 – 4 |
| Vocalization | [0-4] | Vocalization (Normal = 0)<br>The intensity of spontaneous vocalizations can be scored from 0 to 4. |

**S5 Table. Overview of the experimental stage 1 scoring parameters as provided by site 3 and their allocation to the four functional domains for data analysis across sites.**
The results of the parameters allocated to the four functional domains are presented in **Figure 1A, B, C, D**. The results of the parameters subsumed under 'other measures' are presented in **S3 Supplementary Figure**.

| *Parameter* | *Scoring Range* |
|---|---|
| *Motor Domain* | |
| Gait description | [0]-[2] |
| Ataxia | [0]-[2] |
| Posture/ body position | [0]-[2] |
| Righting reflex (air 20 cm) | [0]-[3] |
| Placing reflex | [0]-[1] |
| *Autonomic Domain* | |
| Piloerection | [0]-[1] |
| Lacrimation | [0]-[1] |
| Respiration | [0]-[2] |
| Feces/Diarrhea | [0]-[1] |
| Urination | [0]-[1] |
| Eye prominence | [0]-[1] |
| Salivation | [0]-[2] |
| *Excitation-related Domain* | |
| Stereotypic behavior | [0]-[2] |
| Tremors | [0]-[1] |
| Seizures | [0]-[2] |
| Activity level | [0]-[2] |
| Touch escape | [0]-[2] |
| Startle response | [0]-[2] |
| *Sedation-related Domain* | |
| Position of eyelids | [0]-[2] |
| Activity level | [0]-[2] |
| Touch escape | [0]-[2] |
| Startle response | [0]-[2] |
| *Other Measures* | |
| Tail pinch | [0]-[1] |
| Toe pinch | [0]-[1] |
| General appearance | [0]-[2] |
| Palpebral reflex | [0]-[2] |

**S6 Table. Details about the stage 1 scoring scheme as provided by site 3.**

| *Outcome Measures* | *Scoring Range* | *Description* |
|---|---|---|
| General appearance | [0]-[2] | Normal/unkempt/hunched-moribund |
| Piloerection | [0]-[1] | None; normal/generalized (piloerection all over body) |
| Posture/ body position | [0]-[2] | Normal posture; may include lying down normally/ Extreme excitation – vertical leaping, rearing/ Abnormal – hunched, flattened, laying on side |
| Activity level | [0]-[2] | Normal-active (*active norm) or normal – quiet; resting/ Hyperactivity – extreme movement/ very low arousal/unresponsive |
| Stereotypic behavior | [0]-[2] | Not present/ pild to moderate stereotypy, animals walks out of stereotypic movements/ severe stereotypy, typically interfering with normal behaviors, repetitive interrupted behaviors |
| Gait description | [0]-[2] | Normal/ abnormal slight incapacity/ abnormal moderate to extreme incapacity |
| Ataxia | [0]-[2] | Not present/ moderate overall ataxia as indicated by difficulty moving and coordinating movement/ severe overall ataxia as indicated as inability to move or coordinate movement |
| Eye prominence | [0]-[1] | Not present/present |
| Position of eyelids | [0]-[2] | Open/ ptosis, partially closed lids/ completely closed (*closed) |
| Salivation | [0]-[2] | Not present/ slight: present with narrow area around the mouth involved/ profuse: present with wide area around the mouth and extends to dorsal surface |
| Lacrimation | [0]-[1] | Not present/present |
| Respiration | [0]-[2] | Normal/ abnormal hyper-respiration: rapid breath, chest visibly; abnormal hypo-respiration: dyspnea, difficulty breathing, gasping, labored |
| Tremors | [0]-[1] | Not present / present |
| Seizures | [0]-[2] | Not present / present – mild, transient, pre-seizure activity / present – strong >15 sec in duration with loss of righting reflex, clear behavioral depression after recovery |
| Feces/Diarrhea | [0]-[1] | Not present /present |
| Urination | [0]-[1] | Absent/Present |
| Touch escape | [0]-[2] | Normal – slow escape or slight freeze / Abnormal Hyper response – extremely vigorous, running escape / Abnormal Hypo response - no response |
| Tail pinch | [0]-[1] | Absent/present |
| Toe pinch | [0]-[1] | Absent/present |
| Startle response | [0]-[2] | Normal – slight to moderate reaction/ abnormal hyper response – extremely vigorous / abnormal hypo response - no response |
| Palpebral reflex | [0]-[2] | Normal - blinks (*normal)/ abnormal - no blink (*abnormal) / undetected |
| Placing reflex | [0]-[1] | Normal/abnormal |
| Righting reflex (air 20 cm) | [0]-[3] | Lands on four limbs; normal / all limbs do not touch at once / lands on side / lands on back |

**S7 Table**. **Overview of the experimental stage 1 scoring parameters as provided by site 4 and their allocation to the four functional domains for data analysis across sites.**
The results of the parameters allocated to the four functional domains are presented in **Figure 1A, B, C, D**. The results of the parameters subsumed under 'other measures' are presented in **S4 Supplementary Figure**.

| *Parameter* | *Scoring Range* |
|---|---|
| *Motor Domain* | |
| Abnormal gait | (+)present/(-)absent |
| Akinesia | (+)present/(-)absent |
| Loss of grasping | (+)present/(-)absent |
| Motor incoordination | (+)present/(-)absent |
| Visual orientation loss | (+)present/(-)absent |
| Muscle tone | (+)present/(-)absent |
| *Autonomic Domain* | |
| Piloerection | (+)present/(-)absent |
| Salivation | (+)present/(-)absent |
| Respiratory rate | (+)present/(-)absent |
| Skin colour | (+)present/(-)absent |
| Defaecation | (+)present/(-)absent |
| Ptosis | (+)present/(-)absent |
| *Excitation-related Domain* | |
| Excitation | (+)present/(-)absent |
| Convulsions | (+)present/(-)absent |
| Tremors | (+)present/(-)absent |
| Vocalization | (+)present/(-)absent |
| *Sedation-related Domain* | |
| Sedation | (+)present/(-)absent |
| *Other Measures* | |
| Lethality | (+)present/(-)absent |
| Straub tail | (+)present/(-)absent |
| Wet dog shakes | (+)present/(-)absent |

**S8 Table. Details about the stage 1 scoring scheme as provided by site 4.**

| *Outcome Measures* | *Scoring Range* | *Description* |
|---|---|---|
| Abnormal gait | (+)present/(-)absent | Posture/gait |
| Akinesia | (+)present/(-)absent | Paralysis |
| Convulsions | (+)present/(-)absent | Any convulstion (inspecific) |
| Defaecation | (+)present/(-)absent | Abnormal feces |
| Excitation | (+)present/(-)absent | Increased motoractivity |
| Lethality | (+)present/(-)absent | Death |
| Loss of grasping | (+)present/(-)absent | Mouse is introduced to iron bar which it is supposed to grab when experimenter releases grip off the animal. If it does not "keep hanging in the air", and falls down, this is scored as loss of grasping. |
| Motor incoordination | (+)present/(-)absent | Ataxia |
| Muscle tone | (+)present/(-)absent | Decreased muscle tone |
| Piloerection | (+)present/(-)absent | Piloerection |
| Ptosis | (+)present/(-)absent | Ptosis |
| Respiratory rate | (+)present/(-)absent | Any abnormal respiration (inspecific) |
| Salivation | (+)present/(-)absent | Enhanced salivation |
| Sedation | (+)present/(-)absent | Decreased motoractivity |
| Skin colour | (+)present/(-)absent | Pale, reddened, or blueish |
| Straub tail | (+)present/(-)absent | Straub tail |
| Tremors R - rest tremor | (+)present/(-)absent | Any tremor (inspecific) |
| Visual orientation loss | (+)present/(-)absent | Visual orientation loss (grabbing mouse by tail and slowly moving towards table surface, if it does not reach for the table with front paws before touching the surface this is scored as visual orientation loss) |
| Vocalization | (+)present/(-)absent | Vocalization |
| Wet dog shakes | (+)present/(-)absent | Wet dog shakes |

**S9 Table. Overview of the experimental stage 1 scoring parameters as provided by site 5 and their allocation to the four functional domains for data analysis across sites.**

The results of the parameters allocated to the four functional domains are presented in **Figure 1A, B, C, D**. The results of the parameters subsumed under 'other measures' are presented in **S5 Supplementary Figure**.

| *Parameter* | *Scoring Range* |
|---|---|
| *Motor Domain* | |
| Loss of traction | Presence (+), absence (-) |
| Muscle tone | Increase (+), decrease (-) |
| Abnormal gait | Presence (+), absence (-) |
| Loss of balance | Presence (+), absence (-) |
| Pelvic elevation C - crouched | 0-8 |
| Body position | 0-8 |
| Loss of righting reflex | Presence (+), absence (-) |
| Loss of grasping | Presence (+), absence (-) |
| *Autonomic Domain* | |
| Piloerection | Presence (+), absence (-) |
| Lacrimation | Presence (+), absence (-) |
| Salivation | Presence (+), absence (-) |
| Defaecation | Presence (+), absence (-) |
| Exophthalmia | Presence (+), absence (-) |
| Ptosis | Presence (+), absence (-) |
| *Excitation-related Domain* | |
| Twitches/ convulsions clonic (symmetric/non-symmetric) | 0-8 |
| Twitches/ convulsions tonic | 0-8 |
| Twitches/ convulsions tonic flexion | 0-8 |
| Tremors R - rest tremor | 0-8 |
| Excitation | Presence (+), absence (-) |
| Jumps | Presence (+), absence (-) |
| Snap fingers above cage to evaluate fear | Increase (+), decrease (-) |
| Reactivity to touch | Increase (+), decrease (-) |
| *Sedation-related Domain* | |
| Intensity of sedation | If present: 1 2 3 |
| Snap fingers above cage to evaluate fear | Increase (+), decrease (-) |
| Reactivity to touch | Increase (+), decrease (-) |
| *Other Measures* | |
| Straub tail | Presence (+), absence (-) |
| Loss of corneal reflex | Reflex: presence (+), absence (-) |
| Bizarre behaviour C - circling | 0-8 |
| Lethality | Presence (+), absence (-) |
| Analgesia | Presence (+), absence (-) |
| Aggressiveness towards experimenter | Presence (+), absence (-) |

**S10 Table. Details about the stage 1 scoring scheme as provided by site 5.**

| *Outcome Measures* | *Scoring Range* | *Description* |
|---|---|---|
| Body position | 0-8 | From flat to repeated vertical leaping |
| Intensity of sedation | If present: 1 2 3 | - |
| Jumps | Presence (+), absence (-) | - |
| Sedation | Presence (+), absence (-) | - |
| Straub tail | Presence (+), absence (-) | - |
| Snap fingers above cage to evaluate fear | Increase (+), decrease (-) | From baseline |
| Ptosis | Presence (+), absence (-) | - |
| Exophthalmia | Presence (+), absence (-) | - |
| Reactivity to touch | Increase (+), decrease (-) | From baseline |
| Piloerection | Presence (+), absence (-) | - |
| Loss of grasping | Presence (+), absence (-) | Grasping +/- |
| Loss of corneal reflex | Presence (+), absence (-) | +/- |
| Analgesia | Presence (+), absence (-) | - |
| Loss of traction | Presence (+), absence (-) | Traction +/- |
| Loss of righting reflex | Presence (+), absence (-) | - |
| Lacrimation | Presence (+), absence (-) | - |
| Salivation | Presence (+), absence (-) | - |
| Aggressiveness towards experimenter | Presence (+), absence (-) | - |
| Muscle tone | Increase (+), decrease (-) | - |
| Hypothermia/Hyperthermia | Presence (+), absence (-) - if present: 1 2 3 | - |
| Twitches/ convulsions clonic (symmetric/non-symmetric) | 0-8 | (For increasing magnitude/frequency) |
| Twitches/ convulsions tonic | 0-8 | (For increasing magnitude/frequency) |
| Twitches/convulsions tonic flexion | 0-8 | (For increasing magnitude/frequency) |
| Bizarre behaviour C - circling | 0-8 | - |
| Lethality | Presence (+), absence (-) | - |
| Tremors R - rest tremor | 0-8 | - |
| Defaecation | Presence (+), absence (-) | - |
| Excitation | Presence (+), absence (-) | - |
| Abnormal gait | Presence (+), absence (-) | - |
| Loss of balance | Presence (+), absence (-) | - |
| Pelvic elevation C - crouched | 0-8 | (For flat to markedly elevated) |

**S11 Table. Site-specific variables about animal husbandry, experimental setting, blinding, randomization, and compounds.**

| ***Variable*** | ***Site 1*** | ***Site 2*** | ***Site 3*** | ***Site 4*** | ***Site 5*** |
|---|---|---|---|---|---|
| *Cage type* | Makrolon type III cages (Ehret, Emmendingen, Germany) | Macrolon type II cages | Makrolon type III cages (Ehret, Emmendingen, Germany) | Makrolon type II cages | IVC GR900 solid bottom cages |
| *Temperature and humidity in animal husbandry* | Temperature 20-24℃, Humidity 45-60% | Temperature 20-24°C, Humidity 40-70% | Temperature 20-23°C, Humidity 30-70% | Temperature 20-22℃, Humidity 50-60% | Temperature 21.8-22.5°C, Humidity 35.2-55.3% |
| *Light cycle (...hrs/…hrs) and time of lights on* | 6.15 a.m./ 6.15 p.m. 6.15 am lights on | 6 a.m./6 p.m., 6 a.m. lights on | 7 a.m./ 7 p. m. 7 a.m. lights on | 6 a.m. / 6 p.m. 6 a.m. lights on with 30-minute transition period | 6 a.m. / 6 p.m. 5.30 a.m. lights on |
| *Bedding material* | Aspen chips | Soft wood shavings | Cobs | Aspen chips | Aspen chips |
| *Handling method and frequency* | Twice per week, animals were tail handled with gloved hands and briefly restrained | Tail handling with gloved hands | Twice per week, animals were tail handled with gloved hands and briefly restrained | Once per week tail handled with gloved hands | Once a week during cage change animals were tail handled |
| *Cleaning frequency per week* | Full cage change once a week | Once per week | Full cage change once a week | Full cage change once a week | Full cage change once a week |
| *Water/food replacement frequency, water quality, bedding transfer (yes/no)* | Once a week, tap water, no bedding transfer | Once a week, tap water; bedding transfer (stage 1), no bedding transfer (stage 3) | Once a week, tap water, no bedding transfer | Once a week, tap water, no bedding transfer | Once a week, tap water, bedding transfer yes |
| *Number of care takers interacting with animals* | One | One | Two | One | One |

| *Variable* | *Site 1* | *Site 2* | *Site 3* | *Site 4* | *Site 5* |
|---|---|---|---|---|---|
| *Number of experimenters handling animals* | One | One | Two | One | One |
| *Was experimenter a smoker?* | No | No | No | No | No |
| *Gender of experimenter* | Female | Female | Male | Male | Female |
| *Handling method used by experimenter* | Tail handling with gloved hands | Tail handling with gloved hands | Scruff | Tail handling with gloved hands | Tail handling |
| *Food and/ or water restriction during first testing time points?* | No restrictions | Water and food restriction | No restrictions | Water and food restriction during experiment | No restrictions |
| *Housing during testing period (i.e., between testing time points)* | Home cage | Home cage | Home cage | Home cage | Home cage |
| *Duration of acclimatization to test room* | 1 hour | Unknown | 1 hour | 24 hours - 72 hours | Not reported |
| *Experimental testing arena (home cage or new cage)* | New cage | New cage | New cage | Cylindrical testing "arena" | New cage |
| *Test arena cleaning method* | 0.1 % acetic acid | None | Not provided | None | Water |
| *Test arena bedding* | None | None | Cobs | None | None |
| *Diameter of wire (for testing wire manouvre)* | Outcome measure not recorded | Outcome measure not recorded | Outcome measure not recorded | 3.5 mm | Outcome measure not recorded |
| *Any other relvant details about equipment used for testing time point* | NA | NA | NA | NA | Testing arena was wooden bottom |

| *Variable* | *Stage* | *Site 1* | *Site 2* | *Site 3* | *Site 4* | *Site 5* |
|---|---|---|---|---|---|---|
| *Method applied for randomization of animals (tool/program)* | 1 | The treatments were allocated equally between the sexes (except the group receiving 0.3 mg/kg MK-801, which had one male fewer) using a simple randomization (in-house R script). The order of test cages and order of the animals tested within each cage was randomized (in-house R script, simple randomization). | The animals were randomly put in cages at arrival and assigned to groups cagewise. | The treatments were allocated equally between the sexes using a stratified randomization considering animals' body weight during the week of testing (in-house R script). The order of test cages and order of the animals tested within each cage was randomized (in-house R script, simple randomization). | R-script, simple randomization | Shuffle sealed envelope |
| *Blinding* | 1 | All randomizations were completed by an assistant that did not interact with the animals and the experimenter was blinded to the treatment allocation | Not blinded | All randomizations were completed by an assistant that did not interact with the animals and the experimenter was blinded to the treatment allocation | All randomizations were completed by an assistant that did not interact with the animals and the experimenter was blinded to the treatment allocation | All randomizations were completed by study director that did not interact with the animals and the experimenter was blinded to the treatment allocation |
| *Blinding* | 2, 3 | All randomizations were completed by an assistant that did not interact with the animals and the experimenter was blinded to the treatment allocation | Handler and experimenter were blinded to treatment; animals were dosed by a different person | All randomizations were completed by an assistant that did not interact with the animals and the experimenter was blinded to the treatment allocation | All randomizations were completed by an assistant that did not interact with the animals and the experimenter was blinded to the treatment allocation | All randomizations were completed by study director that did not interact with the animals and the experimenter was blinded to the treatment allocation |

| *Variable* | *Stage* | *Site 1* | *Site 2* | *Site 3* | *Site 4* | *Site 5* |
|---|---|---|---|---|---|---|
| *Provider of vehicle* | 1, 2, 3 | 0.9 % saline, B. Braun, Melsungen, Germany | 0.9 % saline, B. Braun, Melsungen, Germany | In-house | 0.9 % saline, Fresenius Kabi, Germany | 0.9% saline, Baxter Healthcare SA 8010 Zurich, Switzerland |
| *Provider of MK-801* | 1, 2, 3 | Merck, Darmstadt, Germany | Merck, Darmstadt, Germany | In-house | Merck, Darmstadt, Germany | Merck, Darmstadt, Germany |
| *Euthanasia* | 1, 2, 3 | Pentobarbital (Narcoren 16 g/100 ml, 600 mg/kg in 10 ml/kg, i.p.) | Not provided | CO2 asphyxiation | Pentobarbital (Narcoren 16 g/100 ml, 10 ml/kg, i.p.) | CO2 and cervical dislocation |
| *Comments/Deviations from protocol* | 1, 2, 3 | NA | NA | NA | In stage 3, there were deviations from the protocol as the experiments were already scheduled in advance and the stage 3 protocol was finalized too late. Detailed descriptions of the deviations are provided in Supplementary Tables 21 and 22. | No participation in stage 3 |

**S12 Table. Shared stage 2 protocol about harmonization of variables for animal housing and husbandry.**

| ***Variable Factor: Housing and Husbandry*** | ***To be followed*** |
|---|---|
| Number of care takers interacting with animals | Multiple |
| Social housing standardization | Yes – group housed up to single housing 24 hours prior testing. |
| Consistent number per cage | 5 animals per cage if no separation needed due to animals fighting. |
| Environmental enrichment type | Minimum of one enrichment type, which should be documented. |
| Cage standardization | Mouse cage: Makrolon 3108<br>Or albeit to AAALAC guide recommendations which is 6 inches square or 38.7 $cm^2$ for one group housed mouse in a group less than 10 mice |
| Care taker handling method | Tail handling with gloved hands |
| Cleaning frequency per week | Full cage change once a week |
| Water food replacement frequency | Once a week and ad libitum |
| Water quality | Tap water and ad libitum |
| Bedding transfer | None |

**S13 Table. Shared stage 2 protocol about harmonization of variables for experimental animals.**

| ***Variable Factor: Animal*** | ***To be followed*** |
|---|---|
| Strain | NMRI |
| Sex | Male and female |
| Group size | 5/sex/dose |
| Testing age at start | 8 to 8.5 weeks |
| Animal vendor source | Charles River Germany or Taconic Germantown, NY |
| Drug, paradigm and previous procedures naive | Yes |
| Handling habituation frequency | 2 times per week |
| Drink or food restriction pre study | No |

**S14 Table. Shared stage 2 protocol about harmonization of variables for drug treatment and regimen.**

| ***Variable Factor: Drug Treatment and Regimen*** | ***To be followed*** |
|---|---|
| MK-801 | 0; 0.1 and 0.3 mg/kg |
| Route of administration | Intraperitoneal |
| Dose volume | 10 ml/kg |
| Number of dose per animal | One |

**S15 Table. Shared stage 2 protocol about harmonization of variables for drug treatment and regimen.**

| ***Variable Factor Name*** | ***To be followed*** |
|---|---|
| ***Treatment Group Assignments*** | |
| Period of dosing | As per protocol, observation time points are: Baseline, 5, 15, 30, 60, 120, 240 minutes and 24 hrs post dose. |
| Randomization to order animal testing method | R script provided by work package leader |
| Randomization to test group method | Yes |
| Animal characteristics used to balance treatment groups | Body weight |

**S16 Table. Shared stage 2 protocol about harmonization of variables for drug treatment and regimen.**

| ***Variable Name: Personnel*** | ***To be followed*** |
|---|---|
| Experimenter blind during test phase | Yes |
| Handling standardized | Yes |
| Experimenter a smoker | No |

**S17 Table. Shared stage 2 protocol about harmonization of variables for testing set-up.**

| ***Variable Factor Testing set-up*** | ***To be followed*** |
|---|---|
| Time period of experimental assessment | Light phase of dark/light cycle between 2 hours after lights on and 2 hours before lights off. |
| Duration of acclimatization to test room | One hour pre-dosing |
| Experiment environment | Do not use home cage as it should be a new arena |
| Number of animal handlers/experimenter | Single handler |
| Experimenter handling method | Tail handling with gloved hands/gloves |
| Researcher present during test phase | Yes |
| Food restricted during experiment | No food available during the assessment time point |
| Water restriction during experiment | No water available during the assessment time point |
| Housing during testing period | Experimental animals are single housed 24 hours prior first testing time point and remained single house until completion of last testing time point [24 hours post dose] |
| Housed in interval between administration and test | Home cage *in experimental room* |
| Pre-defined humane endpoints | Yes |
| Test area cleaned between animals | Yes |
| Test area cleaning method | 0.1 % acetic acid |
| Test arena bedding type | None |
| Standardization of outcome assessment | Observations by experimenter trained on a fixed assessment protocol (sequence of actions; handling) |

**S18 Table. Shared stage 2 protocol about harmonization of variables for outcome measures.**

| ***Variable Factor: Outcome measures*** | ***To be followed*** |
|---|---|
| Time point assessments | As per protocol, observation time points are: Baseline, 5, 15, 30, 60, 120, 240 minutes and 24hrs post dose. |
| Ex ante primary outcome defined | Each site will not change or add outcomes measured from its original list; however nomenclature and scoring/ rating are harmonized between testing laboratories. |
| Outcome measured scoring system | See table below |
| Ex ante stats analysis plan | Yes |
| Outcomes pre-specified in protocol | Yes |

**S19 Table. Shared stage 2 protocol about harmonization of the report of scoring outcome measures.**

In stage 2, each site continued to use its owned outcome variables panel list which has been used during stage 1. No additional outcomes variables should be added to their own list; however, to help in data comparison, outcome measures nomenclatures and outcome measure rating systems have been harmonized cross sites. Each site should report outcome measures using only nomenclature listed in the first column: "Outcome Measures to be reported" and following the scoring system described in the last column in the table below. The second column reports the different nomenclatures found while reviewing the Localization data and should not be used to report the harmonization data. As an example, if an outcome measure entitled "Gait description" [second column], this should be reported as "Abnormal Gait" [first column] by using the scoring system described on the last column. Body temperature, body weight, glucose level and grip strength will not be incorporated in a cumulative score.

The results of the parameters allocated to the four functional domains are presented in **Figure 2A, B, C, D**. The results of the parameters subsumed under 'other measures' are presented in **S6 Supplementary Figure** (Site 1), **S7 Supplementary Figure** (Site 2), **S8 Supplementary Figure** (Site 3), **S9 Supplementary Figure** (Site 4), and **S10 Supplementary Figure** (Site 5).

| ***Outcome Measures to be reported*** | ***Alternative Outcome Measures Nomenclature*** | ***Scoring Value*** | ***Scoring System*** |
|---|---|---|---|
| Aggression | | [0]-[+1] | Absent =0 or present =+1 |
| Spontaneous activity/ induced activity/ locomotion | | [-1]-[+1] | Slow= -1; active =0; rapid/hyperlocomotion =+1 |
| Convulsions | Seizures | [0]-[+1] | Absent =0 or present =+1 |
| Defecation | Feces/Diarrhea; defecation | [0]-[+1] | Normal* =0 or increased or altered consistency =+1<br>*note that normal can imply absent or present with normal amount or consistency depending on<br>testing situation |
| Piloerection | | [0]-[+1] | Absent =0 or present =+1 |
| Tremors | Temors R - rest tremor; tremor | [0]-[+1] | Absent =0 or present =+1 |
| Visual placing | Placing reflex; visual orientation loss | [0]-[+1] | Present =0 / reduced = +1/ absent =+2 |
| Abnormal gait | Gait description, gait | [0]-[+2] | Normal =0 / abnormal with slight incapacity =+1 / abnormal with moderate to extreme incapacity =+2 |
| Alertness | Sedation; passivity; activity level | [-1]-[+1] | Very low arousal/unresponsive = -1; normal-active (*active norm) or normal – quiet; resting =0 / hyperactivity – extreme movement =+1 |
| Respiration | Respiratory rate L - labored; respiration | [0]-[+2] | Abnormal hypo-respiration: dyspnea, difficulty breathing, gasping, labored =-1 / normal =0 / abnormal hyper-respiration: rapid breath, chest visibly =+1 |
| Salivation | | [0]-[+2] | Not present =0 / slight: present with narrow area around the mouth involved =+1 / profuse: present with wide area around the mouth and extends to dorsal surface =+2 |

| **Outcome Measures to be reported** | **Alternative Outcome Measures Nomenclature** | **Scoring Value** | **Scoring System** |
|---|---|---|---|
| Touch response | Touch escape | [0]-[+2] | Abnormal hypo response - no response =-1 / normal – slow escape or slight freeze =0 / abnormal hyper response – extremely vigorous, running escape =+1 |
| Skin color | General appearance; skin color, C - cyanosis | [0]-[+4] | Pale white = -1 / normal color =0 / light cyanotic =+1; dark cyanotic =+2 |
| Exophthalmoses | Eye prominence, en-/exophthalmus | [0]-[+1] | Absent =0 /present =+1 |
| Irritability | Excitation | [0]-[+1] | Absent =0 /present =+1 |
| Lacrimation | | [0]-[+1] | Normal =0 /increased =+1 |
| Motor incoordination | | [0]-[+1] | Absent =0 /present =+1 |
| Stereotypy | Stereotypic behavior; stereotypies | [0]-[+1] | Absent =0 /present =+1 |
| Straub tail | | [0]-[+1] | Absent =0 /present =+1 |
| Twitches | | [0]-[+1] | Absent =0 /present =+1 |
| Urination | | [0]-[+1] | Normal* =0 /increased amount =+1 *note that normal can imply absent or present with normal amount or consistency depending on testing situation |
| Vocalization | | [0]-[+1] | Absent =0 /present =+1 |
| Wet dog shakes | | [0]-[+1] | Absent =0 /present =+1 |
| Limb position | Posture/ body Position | [0]-[+2] | Normal posture; may include lying down normally =0 /slight abnormality with rotation of hindlimbs = +1/ pronounced abnormality, i.e. flat body posture or lying on side =+2 |
| Palpebral opening | Palpebral closure | [0]-[+2] | Narrowed = -1 / *normal =0 / abnormal, i.e. no blinks and/or widening = +1 |
| Writhing | | [0]-[+1] | absent =0 /present =+1 |
| Ataxia - akinesia | | [0]-[+2] | Not present=0 / moderate overall ataxia as indicated by difficulty moving and coordinating movement =+1 / severe overall ataxia as indicated by inability to move or coordinate movement =+2 |
| Corneal reflex | | [0]-[+2] | Normal = 0 / reduced reflex = +1 / absent, no response = +2 |
| Passivity | | [0]-[+2] | Level of struggle: normal = 0 / reduced = +1 / absent = +2 |
| Alertness/Sedation | | [-1]-[+1] | Sedated = -1 / normal = 0 / pronounced alertness = 1 |
| Activity level | | Normal = 0 ; hyper = +1 ; hypo =-1 | |
| Righting reflex | | [0]-[+2] | Normal =0/ diminished =-1 / completely absent =-2 |

| Outcome Measures to be reported | Alternative Outcome Measures Nomenclature | Scoring Value | Scoring System |
|---|---|---|---|
| Toe pinch reflex | Toe pinch; tail pinch, pinna reflex | [0]-[+2] | Normal =0/ diminished = +1 /absent = +2 |
| Wire maneuver | Grip strength; loss of grasping | [0]-[+2] | Normal =0/diminished = +1/absent = +2 |
| Catalepsy | | [0]-[+1] | Catalepsy (normal = 0) /animal is completely cataleptic =+1 |
| Extensor reflex | | [0]-[+2] | Normal = 0 / reduced = +1 /absent, no response = +2 |
| Flexor reflex | | [0]-[+2] | Normal = 0 / reduced = +1 /absent, no response = +2 |
| Grooming | | [-1]-[+1] | Reduced grooming behavior or no grooming = -1/normal grooming =0/ increased grooming behavior = +1 |
| Pain response | | [-1]-[+1] | No pain reaction = -1/ normal response =0/ high pain response =+1 |
| Pupil size | | [-1]-[+1] | Pupil shrinking =-1 / normal pupil size =0/ dilation of pupil =+1 |
| Struggle response | | [-1]-[+1] | Normal reaction to being scruffed =0; animal lies calmly on the hand and easy to scruff =-1; animal struggles =+1 |
| Fearfulness | | [-1]-[+1] | The investigator slowly approaches mouse with hand (before picking up to transfer). Animal approaches hand = -1 / normal =0 / pronounced freezing/vigorous escape = +1 |
| Pupil size | | [-1]-[+1] | Miosis/constricted =-1 / normal =0 / mydriasis/dilated =+1 |
| Startle response | | [-1]-[+1] | Diminished =-1 / normal =0 / exaggerated response =+1 |
| Body temperature | Rectal temperature | Value | Record the value shown on the device. |
| Body weight | | Value | Record the value shown on the device. |
| Glucose level | | Value | Record the value shown on the device. |
| Grip strength | Wire maneuver; loss of grasping | Value | Record the value shown on the device. |

**S20 Table. Overview of the experimental stage 3 scoring parameters as provided by the shared protocol.** This shared protocol was followed by site 1, 2, and 3. The overview indicates the allocation of the parameters to the four functional domains for centralized data analysis across sites. The results of the parameters allocated to the four functional domains are presented in **Figure 3A, B, C, D**. The results of the parameters subsumed under 'other measures' are presented in **S11 Supplementary Figure**.

*The parameters pupil size and increased/decreased startle response were not included in the functional domains for centralized analysis. Due to the very low light level during experiments at site 1 (which had to be maintained as it was used in previous stages), pupil size was not possible to judge and this parameter was therefore removed from centralized analysis.

| *Parameter* | *Scoring Range* |
|---|---|
| *Motor Domain* | |
| Wire manouvre | 0, 1 |
| Visual placing | 0, 1 |
| Abnormla gait | 0, 1, 2 |
| Limb position | 0, 1 |
| Righting reflex | 0, 1 |
| Motor incoordination | 0, 1, 2 |
| *Autonomic Domain* | |
| Urination | 0, 1 |
| Defecation | 0, 1 |
| Respiration | 0, 1, 2 |
| Skin color | 0, 1 |
| *Pupil size | 0, 1 |
| Lacrimation | 0, 1 |
| *Excitation-related Domain* | |
| Tremors | 0, 1 |
| Convulsions | 0, 1 |
| Vocalization | 0, 1 |
| Hyperactivity | 0, 1, 2 |
| Stereotypy | 0, 1 |
| Increased touch response | 0, 1 |
| Increased irritability | 0, 1 |
| Increased fearfulness | 0, 1 |
| Increased alertness | 0, 1 |
| *Sedation-related Domain* | |
| Hypoactivity | 0, 1, 2 |
| Decreased touch response | 0, 1 |
| Decreased irritability | 0, 1 |
| Decreased fearfulness | 0, 1 |
| Decreased alertness | 0, 1 |
| *Other Measures* | |
| Writhing | 0, 1 |
| Loss of tail/toe pinch reflex | 0, 1 |
| Loss of corneal reflex | 0, 1 |
| Startle response | 0, 1 |

**S21 Table. Overview of the experimental stage 3 scoring parameters used by lab 4.**
In deviation from the shared protocol, the scoring scheme provided below was only used by lab 4 in stage 3. The overview indicates the allocation of the parameters to the four functional domains for analysis.
* The parameter 'wire manouvre' was excluded from analysis due to missing values from male animals.
The results of the parameters allocated to the four functional domains are presented in **S12 Supplementary Figure**. The results of the parameters subsumed under 'other measures' are presented in **S13 Supplementary Figure**.

| *Parameter* | *Scoring Range* |
|---|---|
| *Motor Domain* | |
| *Wire manouvre | 0, 1 |
| Visual placing | 0, 1 |
| Abnormal gait | 0, 1, 2 |
| Limb position | 0, 1 |
| Righting reflex | 0, 1 |
| Motor incoordination | 0, 1, 2 |
| Decreased muscle tone | 0, 1 |
| *Autonomic Domain* | |
| Respiration | 0, 1, 2 |
| Skin color | 0, 1 |
| Lacrimation | 0, 1 |
| *Excitation-related Domain* | |
| Tremors | 0, 1 |
| Convulsions | 0, 1 |
| Vocalization | 0, 1 |
| Hyperactivity | 0, 1, 2 |
| Stereotypy | 0, 1 |
| Increased alertness | 0, 1 |
| *Sedation-related Domain* | |
| Hypoactivity | 0, 1, 2 |
| Decreased alertness | 0, 1 |
| *Other Measures* | |
| Startle respsone | 0,1 |
| Writhing | 0, 1 |

**S22 Table. Overview of the experimental stage 3 scoring parameters which were not measured by lab 4.** These parameters are given in the shared stage 3 protocol, but were not recorded by lab 4.

| *Missing Outcome Measure* | *Scoring Range* |
|---|---|
| Urination | 0, 1 |
| Defecation | 0, 1 |
| Pupil size | 0, 1 |
| Increased touch response | 0, 1 |
| Increased irritability | 0, 1 |
| Increased fearfulness | 0, 1 |
| Decreased touch response | 0, 1 |
| Decreased irritability | 0, 1 |
| Decreased fearfulness | 0, 1 |
| Tail/toe pinch reflex | 0, 1 |
| Corneal reflex | 0, 1 |
